# Blm10 scaffolds 14-3-3 adaptors to the proteasome, and maintains cellular proteostatic balance and gamete quality

**DOI:** 10.64898/2025.12.08.693079

**Authors:** Maia C. Reyes, Diego R. Ramos-Ortiz, Jiarong A. Cheng, Silvan Spiri, Ella Doron-Mandel, Jenny Kim Kim, Yanzhe Ma, Marko Jovanovic, Andreas Martin, Gloria A. Brar

**Author notes:** These authors contributed equally (Maia C. Reyes, Diego R. Ramos-Ortiz).

## Abstract

The proteasome is responsible for regulated protein degradation in eukaryotic cells. Its best characterized substrates are ubiquitinated proteins that are targeted to the 26S proteasome complex, consisting of a 19S regulatory particle (RP) capping the barrel-shaped 20S core peptidase (CP). The CP can interact with other caps, including Blm10/PA200, a nearly 250 kDa protein whose biological function is not well understood. Blm10 is upregulated during gametogenesis in budding yeast, suggestive of a natural stage-specific modulation of proteasome composition. Here, we investigate the function of Blm10 during yeast gametogenesis, identifying it as a weak activator of the proteasome that can displace the 19S RP from the CP. Due to this competition for the CP, overexpression of Blm10 can lead to attenuation of ubiquitin-dependent degradation and proteostatic defects. Cells lacking Blm10 also display markers of proteostatic stress, including Hsp104 foci and heat sensitivity, suggesting that Blm10 safeguards normal proteostatic balance. We find that Blm10 is important for producing fit gametes and ensuring full rejuvenation of aged cells following gametogenesis. Furthermore, we observe direct association of aggregate-prone proteins and protein-folding factors with immunoprecipitated Blm10-proteasomes. Finally, we discovered that 14-3-3 proteins Bmh1/2 are novel cofactors of Blm-10 bound proteasomes and describe how Blm10 controls CP gate configuration. Overall, our data suggest a role for Blm10-proteasomes in maintaining gamete proteostasis through fine-tuning of proteasome activity and prevention of protein aggregation.

## Introduction

Proteostasis, the maintenance of protein homeostasis, is essential for cellular fitness. Perturbations to the proteolytic machinery and a decline in proteostasis are associated with aging and disease (Hipp et al., 2019; Klaips et al., 2017; Labbadia & Morimoto, 2015). The 26S proteosome is a central regulator of protein degradation, a key mediator of proteostasis, and it has been most studied for its role in the selective degradation of ubiquitin-labeled proteins in an ATP-dependent manner (Ciechanover, 1998; Dikic, 2017; Kandel et al., 2024). These protein targets include damaged, misfolded, and “unwanted” but otherwise functional proteins that are not required by the cell in a specific context (Ciechanover, 1998; Dikic, 2017; Kandel et al., 2024).

The 26S proteasome is composed of two major subcomplexes. The 20S core peptidase (CP) contains two outer and two inner rings of α and β subunits that form a proteolytic barrel with sequestered active sites (Budenholzer et al., 2017; Marshall & Vierstra, 2019; Tanaka, 2009). The outer α-rings include a gate that limits access to three proteolytic subunits within the inner β-ring (with caspase-, trypsin-, and chymotrypsin-like activity) which are ultimately responsible for the degradation of the target protein (Budenholzer et al., 2017; Marshall & Vierstra, 2019; Tanaka, 2009). The 19S regulatory particle (RP) caps one or both apical ends of the 20S CP, captures ubiquitinated substrates, and uses its six ATPase subunits of the AAA+ family to mechanically unfold and translocate substrates into the CP for degradation (Arkinson et al., 2025; Bard et al., 2018; Budenholzer et al., 2017; Marshall & Vierstra, 2019; Schmidt & Finley, 2014; Tanaka, 2009). Some of these ATPase subunits also contain HbYX motifs in their C-terminal tails that dock into hydrophobic pockets of the CP α-ring and facilitate gate opening for substrate entry into the degradation chamber (Arkinson et al., 2025; Bard et al., 2018; Budenholzer et al., 2017; Marshall & Vierstra, 2019; Schmidt & Finley, 2014; Tanaka, 2009).

The activity and function of the proteasome can be modulated by modification of its composition. Specialized proteasomes with unique CP composition, including immunoproteasomes, thymoproteasomes, and spermatoproteasomes have been identified (Abi Habib et al., 2022; Gómez-H et al., 2019; Morozov & Karpov, 2019; Murata et al., 2018; Xiong et al., 2022). Additionally, the CP can interact with “caps” other than the 19S RP to modulate function of the overall complex, including the conserved regulator Blm10/PA200 (Schmidt et al., 2005). Blm10 is a large, dome-shaped protein primarily composed of solenoid loops of HEAT-like repeats (Iwanczyk et al., 2006; Sadre-Bazzaz et al., 2010; Schmidt et al., 2005). The function of Blm10-bound proteasomes is not well understood but possibly involves ubiquitin-and ATP-independent substrate degradation. Previous studies have implicated Blm10 in diverse roles including the involvement in DNA repair (Moore, 1991; Ustrell et al., 2002), nuclear import of the proteasome (Weberruss et al., 2013), proteasome maturation and assembly (Fehlker et al., 2003; Kaur et al., 2024), aging and epigenetic maintenance (L.-B. Chen et al., 2020; Y.-S. Chen et al., 2021; Qian et al., 2013), and murine spermatogenesis (Huang et al., 2016; Khor et al., 2006; Sato et al., 2023). Previous studies have also reported select targets of Blm10-proteasomes, including mitochondrial fission protein Dnm1 (Tar et al., 2014), ribosome transcription factor Sfp1 (Lopez et al., 2011), tau fibrils *in vitro* (Dange et al., 2011), N-terminal huntingtin fragments (Aladdin et al., 2020), and acetylated histones (Y.-S. Chen et al., 2021; Qian et al., 2013). Based on the lack of a large pore in the Blm10 cap and its absence of ATPase activity, it has been suggested that substrates of Blm10-proteasomes may be small peptides or already unstructured proteins that do not require ATP-dependent unfolding and translocation into the CP for degradation (Dange et al., 2011).

Blm10 is upregulated during gametogenesis in budding yeast, the conserved process by which diploid cells differentiate into haploid gametes (Eisenberg et al., 2018). This is a highly regulated process during which precise waves of gene regulation ensure that cells can segregate cellular components including chromatin, organelles, and proteins, in a concerted and timely manner to generate gametes (Brar et al., 2012; Marston & Amon, 2004; Neiman, 2011). Additionally, gametogenesis benefits from quality control mechanisms to ensure that gametes contain high-quality components (Sing et al., 2022; Ünal et al., 2011; Xiao & Ünal, 2025). For example, protein aggregates are excluded from inheritance into gametes and eliminated (King et al., 2019; Ünal et al., 2011). The potency of these quality control mechanisms is reflected in the remarkable ability of gametogenesis to reset the replicative lifespan of all four gametes produced from aged precursor cells (Ünal et al., 2011).

In this study, we investigate the function of Blm10-proteasomes during gametogenesis. We find that Blm10-proteasomes are enriched during late gametogenesis and are catalytically active in cleaving unstructured peptides, but less efficient at doing so than the 26S proteasome. Interestingly, we found that either too little or too much Blm10 disrupts normal cellular proteostasis, leading to accumulation of aggregated proteins, and that Blm10 is important for generating high-quality gametes and rejuvenating the gametes of aged precursor cells. Finally, we describe how the conformational landscape of Blm10-bound proteasomes controls CP gate access and report stable association of Blm10-proteasomes with 14-3-3 proteins, which we propose act as cofactors to determine the targets of this non-canonical proteasome assembly.

## Results

### Blm10-proteasome complexes are enriched during late gametogenesis

The RP is the most studied cap of the proteasome, but it is known that Blm10 can also bind in a mutually exclusive manner to either side of the CP in budding yeast. Proteasome assemblies with a combination of one, two, or no caps have been observed (Figure 1A, (Schmidt et al., 2005)). In a previous study, we leveraged parallel global measurements of mRNA, translation, and protein abundance for cells during gametogenesis to analyze protein complex regulation (Cheng et al., 2018; Eisenberg et al., 2018). We observed that CP and RP subunits appear to be co-regulated at the level of mRNA abundance and translation, consistent with their transcriptional induction by the shared transcriptional regulator Rpn4 (Figure 1B, (Mannhaupt et al., 1999)). However, at the protein level, RP subunits decrease in abundance during late gametogenesis, from about 8 to 24 hours in sporulation media (“SPO”), a time period corresponding with the events of late Meiosis II and gamete maturation (gray box, Figure 1B). During this window, Blm10 mRNA, translation, and protein increase dramatically (gray box, Figure 1B) suggesting a stage-specific modulation in proteasome composition whereby Blm10 may be replacing RPs on a subset of CPs.

**Figure 1.**
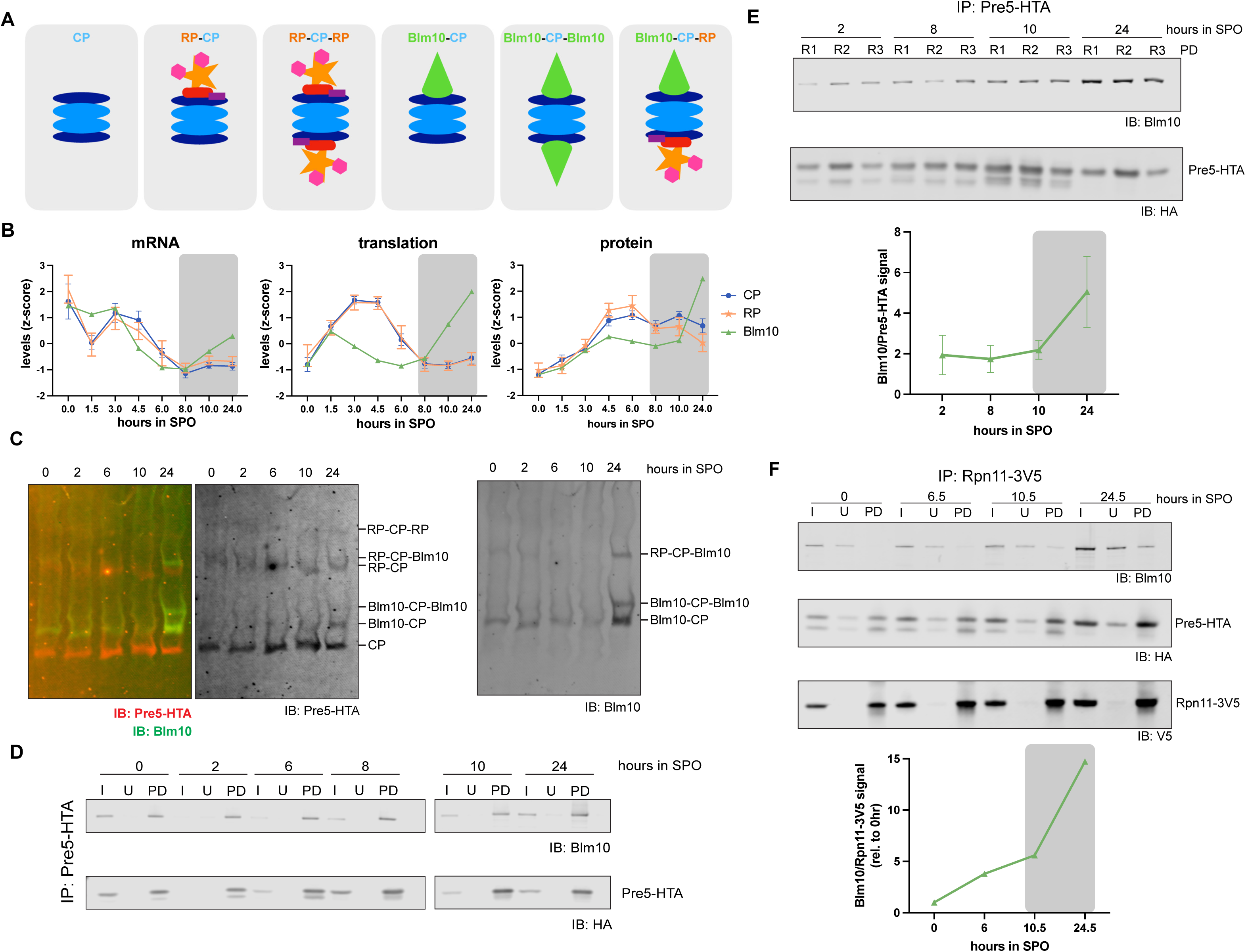
Blm10-proteasomes are enriched during late gametogenesis. (A) Schematic of the major proteasome complexes in budding yeast. Components include the 20S core peptidase (“CP,” blue), 19S regulatory particle (“RP,” orange), and Blm10 (green). (B) mRNA, translation, and protein-level measurements of CP, RP, and Blm10 throughout gametogenesis (adapted from (Eisenberg et al., 2018)). Blm10 is upregulated during late gametogenesis (gray box, ∼8-24 hours in sporulation media/SPO). (C) Native immunoblot of meiotic lysates for Pre5-HTA (CP protein) and Blm10. Replicate in Figure S1A. (D) Co-IP of Pre5-HTA across indicated timepoints in SPO. I = input, U = unbound, PD = pulldown. (E) Co-IP of Pre5-HTA across a separate meiotic timecourse in SPO. Pulldown fractions in triplicate (R1, R2, R3) are shown. Quantification of the amount of Blm10 that co-immunoprecipitated with Pre5-HTA is shown below, with increase from 10-24 hours in SPO (gray box). (F) Co-IP of Rpn11-3V5 (RP protein). I = input, U = unbound, PD = pulldown. Quantification of the amount of Blm10 that co-immunoprecipitated with Rpn11-3V5 is shown below, with increase from 10.5-24.5 hours in SPO (gray box).

We confirmed the increase in Blm10 association with proteasomes during late gametogenesis via resolving proteasome complexes from meiotic cell lysates on native gels, and immunoblotting for Blm10 and Pre5-HTA, a tagged CP protein (Figure 1C). Proteasome complexes run with a characteristic banding pattern on native gels (Roelofs et al., 2018). Consistent with the global measurements, Blm10-proteasomes are highly abundant at 24 hours in SPO (Figure 1C). We additionally confirmed the increase in Blm10-proteasomes by performing co-immunoprecipitations (co-IPs) with Pre5-HTA from meiotic lysates (Figure 1D, 1E). At 24 hours in SPO, more Blm10 co-immunoprecipitated with the CP than at earlier timepoints (Figure 1E). Additionally, we observed that virtually all Blm10 in our cell lysate was bound to the CP, based on the absence of Blm10 in the unbound fraction of co-IPs and the absence of a free Blm10 band in the native immunoblots (Figure 1C, 1D, 1E, S1A (Burris et al., 2021)). Moreover, mass spectrometry data of cell extracts show that abundance of Blm10 is on par with that of RP components in cells at 24 hours in SPO but not an earlier timepoint (Figure S1B). Mass spectrometry of immunoprecipitated CPs from 24 hours in SPO revealed Blm10 to be one of the top CP interactors in gametes, with levels of association rivaling RP components (Figure S1C). Using gel-based assays, we could not reproducibly detect the decrease in RP-CP-RP or RP-CP predicted by our proteomic measurements (Figure 1B, 1C, S1A) in late meiotic lysates. This may be due to the difference in sensitivity between these assays, as well as the difficulty in maintaining physiological RP-CP interactions, which are ATP-dependent (Liu et al., 2006), upon cell lysis. However, we did observe that the abundance of hybrid RP-CP-Blm10 structures increases steadily during meiosis and is highest after 24 hours, with similar timing to Blm10 upregulation and Blm10-CP/Blm10-CP-Blm10 population (Figure 1F).

### Blm10-proteasomes cleave a fluorogenic peptide substrate less efficiently than 26S species

We sought to further characterize the properties of Blm10-proteasomes in our system using a biochemical approach. Using a β-estradiol (β-ER) -inducible LexA trans-activator to drive *BLM10*expression under the control of upstream LexO sites (Figure 2A, (Ottoz et al., 2014)), we collected mitotic cell lysates to analyze proteasomal peptide-cleavage activity via an in-gel native peptidase activity assay. Proteasome complexes are visualized by incubating the gel with a reaction mixture containing the fluorogenic substrate Succinyl-LeuLeuValTyr-7-Amino-4-methylcoumarin (Suc-LLVY-AMC), which is a specific substrate for the chymotrypsin-like proteolytic site in the CP (Roelofs et al., 2018). Catalytically stimulated proteasomes, i.e. ones whose 20S CP gates are opened by the RP for substrate entry, cleave after the LLVY sequence and thereby release and unquench the fluorescent AMC moiety, causing the associated band to be visible (Figure 2B, (Roelofs et al., 2018)). Only with the addition of low concentrations of SDS (0.02%), which is known to artificially open the gates of the auto-inhibited CP (Roelofs et al., 2018), were we able to visualize activity for the Blm10-bound proteasomes (Figure 2B). Blm10 has a single C-terminal HbYX motif, while other characterized proteasome activators such as the RP have several of these motifs to dock and activate the CP (Dange et al., 2011; Sadre-Bazzaz et al., 2010). The requirement of SDS addition to visualize Blm10-proteasomes on native gels is consistent with some previous studies (Lehmann et al., 2008; Sahoo et al., 2024). This suggests that the degradative activity of Blm10-proteasomes is lower than that of 26S proteasomes, likely due to a more restricted substrate access to the degradation chamber. We confirmed the formation of Blm10-proteasomes in this inducible system via native immunoblot (Figure 2C). We also observed that the activity of Blm10-proteasomes in meiotic lysates also require 0.02% SDS to be visualized on native gels as well (Figure 2D). Blm10-CP-Blm10 proteasomes in meiotic lysates were not resolved in this assay, as the amounts of Blm10-CP-Blm10 in meiotic lysates may be too low for detection (Figure 1C, immunoblot).

**Figure 2.**
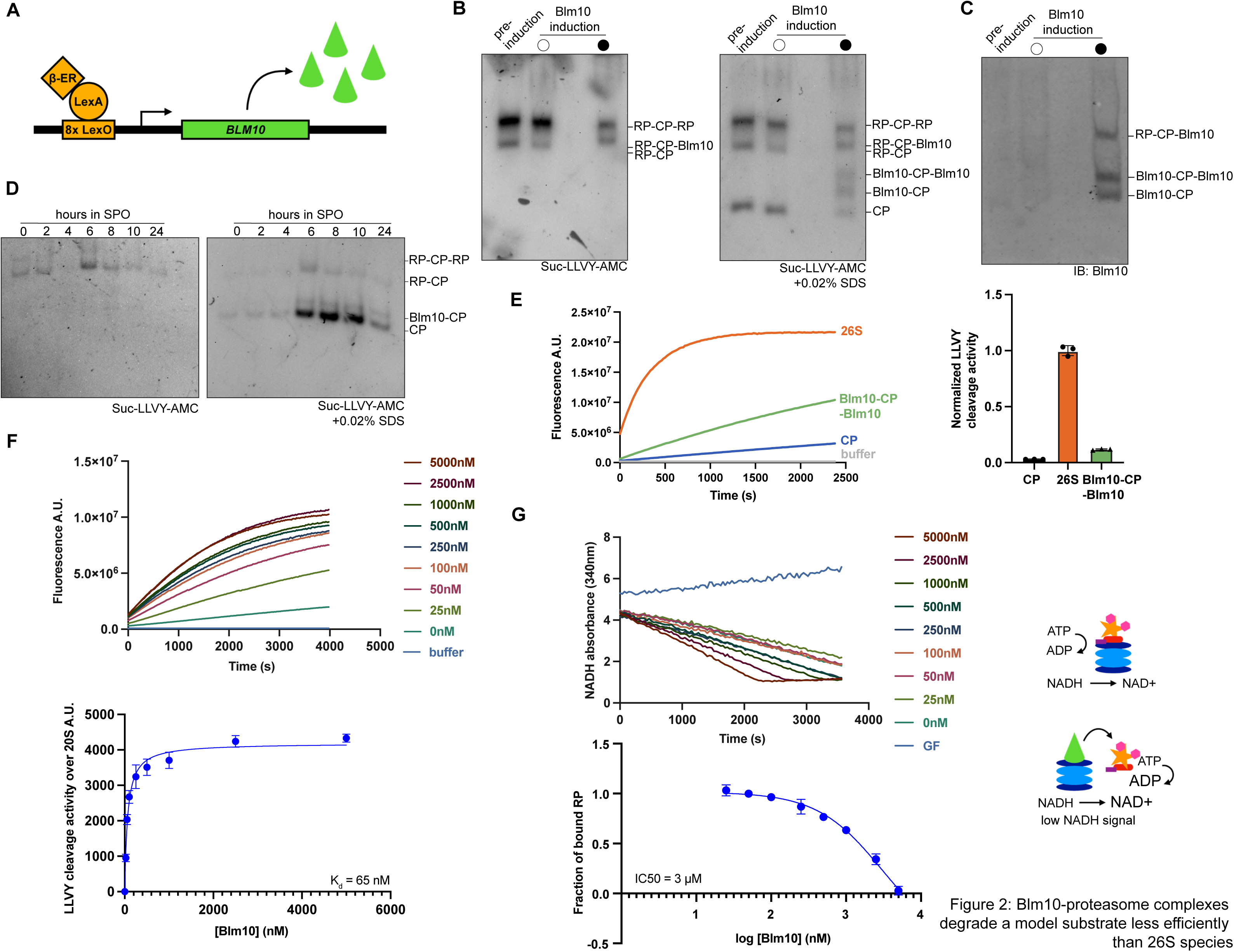
Blm10-proteasome complexes have weak catalytic activity. (A) Schematic of the beta-estradiol (β-ER) inducible LexA/LexO system used to overexpress *BLM10* (Ottoz et al., 2014). LexA, a trans-activator that also contains an estrogen receptor (ER) domain, stimulates the expression of genes downstream of LexO sites upon the addition of 1µM β-ER. This system is used in cells with the endogenous *BLM10* locus deleted. (B) Native gel peptidase assay of inducible Blm10 samples, using the fluorogenic substrate Suc-LLVY-AMC, which is a specific substrate for the chymotryptic-like proteolytic site of the proteasome. Upon addition of 0.02% SDS to the reaction mixture, the activity of the CP, Blm10-CP-Blm10, and Blm10-CP complexes are visualized (right). (C) Native immunoblot for Blm10, confirming the formation of Blm10-containing proteasomes after Blm10 induction. (D) Native gel peptidase assay, using Suc-LLVY-AMC, of meiotic lysates from indicated timepoints in SPO. The activity of CP and Blm10-CP complexes is visualized with the addition of 0.02% SDS (right). (E) Representative LLVY cleavage activity measured for 25 nM of the following purified complexes from one replicate: 26S proteasome (orange), Blm10-proteasomes (green), CP (blue), and buffer (gray). Measurements are for assumed doubly capped 26S and doubly capped Blm10-proteasomes. Triplicate data for normalized LLVY cleavage activity (relative to the 26S) is shown to the right. (F) Representative LLVY cleavage activity generated from increasing purified [3x-Flag-Blm10] mixed with 25 nM purified CP from one replicate. Triplicate data fitted to binding curve (K_d_ = 65 nM, below). Native immunoblot of select concentrations of Blm10 mixed with purified CP in Figure S2A. (G) Representative ATPase displacement assay for measurement of RP bound to CP in purified 100 nM 26S proteasomes, in the presence of increasing [3x-Flag-Blm10], for one replicate. NADH oxidation is coupled to the amount of ATP that is consumed by the RP. NADH absorbance (at 340 nm) is shown over time. Triplicate data fitted to displacement curve (IC50 = 3 μM, below). Native immunoblot of select samples in Figure S2B, S2C.

To further characterize Blm10-proteasomes, we performed *in vitro* analysis. We compared the catalytic activity of purified complexes by measuring the increase in AMC fluorescence over time during the peptidase assay (Figure 2E). As expected, 26S proteasomes displayed the highest peptidase activity, and the isolated CP displayed a low level of basal peptidase activity (Figure 2E). The activity of Blm10-CP complexes exceeds the basal CP level but is low compared to the 26S proteasome, consistent with our in-gel assay results (Figures 2B, 2E).

### Blm10 competes with RP for CP binding

We noted that induction of Blm10 expression reduced the level of free CP and also RP-CP species in our native gel analysis, suggesting that it can effectively compete *in vivo* with RP for CP binding, even when free CP is present (Figure 2B). To investigate the binding affinity of Blm10 to the CP, purified 3x-Flag-tagged Blm10 was titrated into a fixed concentration of purified CP (Figure 2F). Since Blm10-bound proteasomes display catalytic activity, the Suc-LLVY-AMC peptidase assay was used as a proxy for binding. We determined that Blm10 binds to the CP with a Kd of ∼65 nM (Figure 2F). In addition, by mixing varying concentrations of Blm10 with CP and performing immunoblotting after native PAGE we confirmed that the expected proteasome complexes (Blm10-CP-Blm10, Blm10-CP) were formed (Figure S2A).

We next investigated whether Blm10 was capable of competing with RP for CP binding. Purified 3x-Flag-Blm10 was titrated into a fixed concentration of purified 26S proteasome, and an established ATPase assay was used to estimate RP dissociation (Figure 2G). In this assay, NADH oxidation is coupled to ATP consumption, and it has been previously shown that CP binding reduces the RP’s ATPase activity by ∼ 2-fold, with a K_d_ of 127 nM (Beckwith et al., 2013)). With increasing Blm10 concentration, we observed an apparently complete RP displacement from CP (Figure 2G, IC50 = 3 μM). This is in agreement with a previous study that observed reduction of the 26S proteasome upon strong overexpression of *BLM10* in budding yeast lysates (Burris et al., 2021). By mixing increasing concentrations of Blm10 with a fixed amount of 26S proteasome and performing immunoblotting after native PAGE, we again confirmed that the expected proteasome species (Blm10-CP, Blm10-CP-RP, Blm10-CP-Blm10) were forming (Figure S2B, S2C). Notably, we also observed the appearance of unbound RP, signifying its CP displacement by Blm10 (Figure S2B).

### Loss of Blm10 results in minimal disruption of the meiotic proteome

To determine the global effects of Blm10 on gene expression during gametogenesis, we analyzed protein and mRNA levels in parallel across a meiotic time course by label free mass spectrometry and matched mRNA-seq of extracts from WT and *blm10Δ* cells (Figure 3A, 3B respectively). The data were reproducible, as judged by analysis of replicate samples (Figure S3). Few differences were observed, consistent with the normal meiotic progression of cells lacking Blm10. However, several subtle differences could be seen, including two clusters of proteins that were higher at late timepoints in cells lacking Blm10 relative to WT controls and that were enriched for mitochondrial proteins and catabolic roles (Figure 3A, “slow decline 1” and “slow decline 2”). Comparison to mRNA-seq data indicated that most of these changes were also seen at the level of mRNAs, suggesting that they may reflect secondary cellular responses to the lack of Blm10 rather than changes in protein stability mediated by Blm10-bound proteasomes (Figure 3C). This conclusion was made based on analyses of the correlation between mRNA and protein levels over the time course of the experiment in WT and *blm10Δ* cells, showing generally similar mRNA and protein level changes for these genes over the time course (Figure 3C).

**Figure 3.**
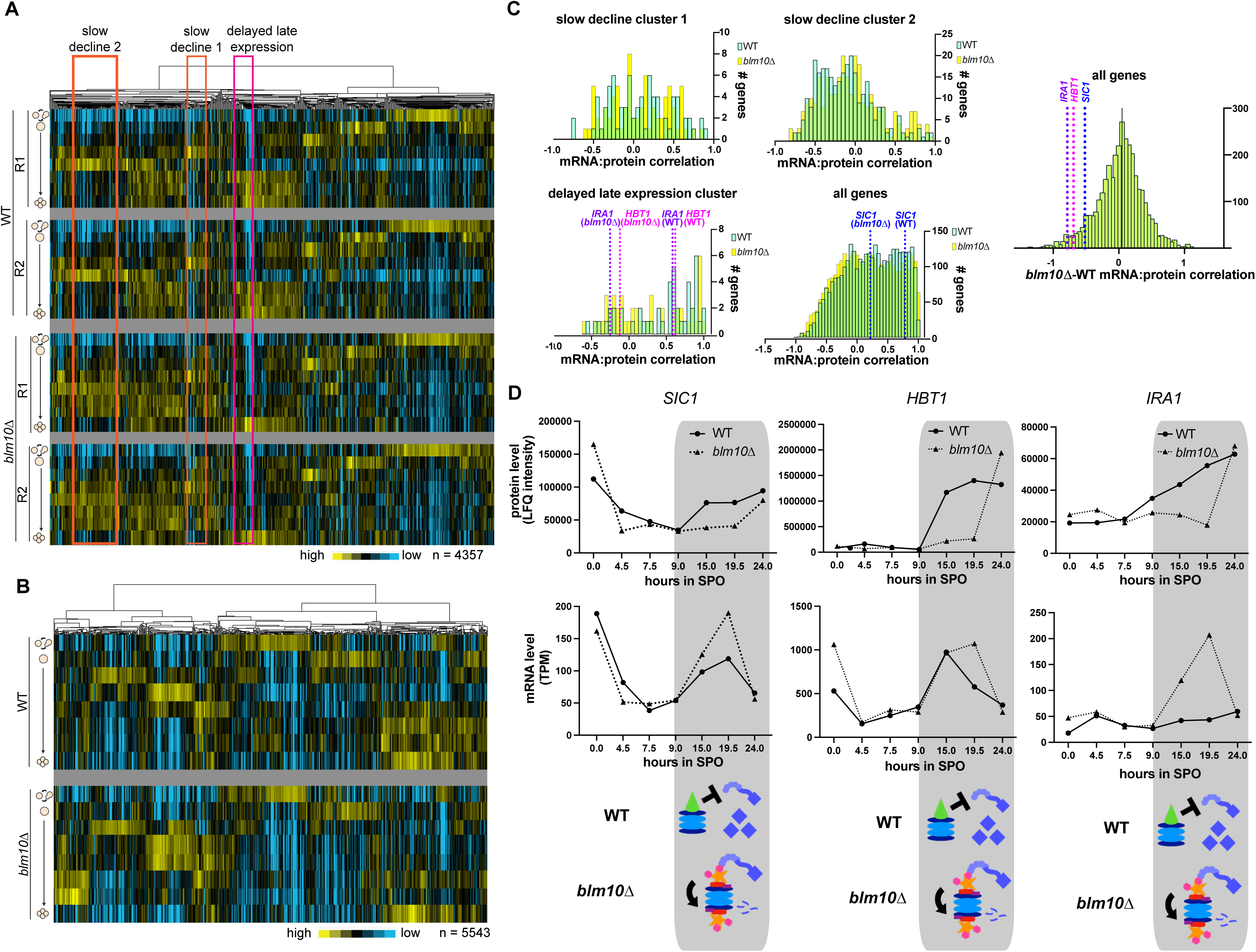
26S proteasome targets are hyperdegraded in *blm10*Δ cells during gametogenesis. (A) Global view of mass spectrometry (MS) data (label-free quantification (LFQ) intensity, normalized across genes and hierarchical clustered) for WT and *blm10*Δ cells throughout gametogenesis. Each row is a timepoint, starting with a mitotic sample, followed by meiotic timepoints. R1 = replicate 1, R2 = replicate 2. N = 4357 proteins identified. (B) Global view of matched mRNA-seq data (transcript per million (TPM), normalized across genes and hierarchical clustered), for WT and *blm10*Δ cells throughout meiosis. Each row is a timepoint, starting with a mitotic sample, followed by meiotic timepoints. N = 5543 transcripts identified. (C) Histograms of clusters shown in (A) (slow decline cluster 1, slow decline cluster 2, delayed late expression cluster) in WT (light blue) or *blm10*Δ (yellow) mRNA:protein correlation data. Overlap in the number of genes per bin between WT and *blm10*Δ appears as green. In the “delayed late expression” cluster, 26S proteasome targets Ira (purple) and Hbt1 (pink) have poorer mRNA:protein correlation in *blm10*Δ cells. When examining all genes, 26S proteasome target Sic1 (dark blue) is shown as another example of a gene with poorer mRNA:protein correlation in *blm10*Δ cells. (D) Protein levels (LFQ intensity) and mRNA levels (TPM) for 26S targets Sic1, Hbt1, and Ira1 in WT and *blm10*Δ meiotic timecourses. Gray box is the timeframe of Blm10 upregulation during gametogenesis. Model for hyperdegradation of 26S targets is shown below. In WT cells, Blm10-proteasomes are present and Sic1, Hbt1, and Ira1 accumulate. In *blm10*Δ cells, decreased levels of Sic1, Hbt1, and Ira1 are observed, suggesting that hyperdegradation by the 26S proteasome is occurring. mRNA levels of Sic1, Hbt1, and Ira1 also increase in *blm10*Δ cells, suggesting feedback.

### 26S proteasome targets are hyperdegraded in *blm10*Δ cells

Interestingly, for cells lacking Blm10 we also observed a delay in the expression of a group of genes enriched for roles in cellular detoxification in cells lacking Blm10 (Figure 3A, “delayed late expression”). Some of these genes displayed a poorer correlation between mRNA and protein trends in this time course, when comparing WT and *blm10*Δ samples (Figure 3C). Two genes that showed an especially large Blm10-dependent discordance between mRNA and protein abundance were *IRA1*/neurofibromin and *HBT1* (Figure 3C, 3D). Strikingly, both Ira1 and Hbt1 are known to be subject to 26S proteasome-mediated degradation (Cichowski et al., 2003; Fang et al., 2011). Intriguingly, we also noted that in the absence of Blm10, another known 26S proteasome target, Sic1 (Verma et al., 2001), shows delayed protein accumulation during late gametogenesis (Figure 3D). The timed degradation and buildup of Sic1 protein is important for cell cycling, and in *blm10Δ* cells the re-accumulation of Sic1 protein is delayed by about 4.5 hours, in a timeframe in which *BLM10* is upregulated (Figure 3D). Similar to Ira1 and Hbt1, Sic1 mRNA levels actually increase in cells lacking Blm10 indicating potential feedback as a result of hyperdegradation (Figure 3D). These findings are consistent with a model in which the presence of high Blm10 concentrations late in meiosis normally compete with RPs for CP binding, and thus 26S proteasome activity is enhanced in Blm10’s absence. Because Blm10-proteasomes do not degrade ubiquitinated substrates, and the RP can bind the CP without competition in *blm10*Δ cells, Sic1, Ira1, and Hbt1 are hyperdegraded, whereas in gametes with high Blm10 abundance, these proteins accumulate normally (Figure 3D).

### Either high or low Blm10 abundance disrupts proteostasis

It is known that dysregulation of the ubiquitin-proteasome system can lead to a decline in proteostasis (Klaips et al., 2017; Labbadia & Morimoto, 2015). To further investigate the physiological relevance of Blm10 for cell health, we examined the impact of Blm10 levels using overexpression studies. In either the absence of the Tet repressor (TetR) (Figure 4A, Figure S4A) or with the addition of anhydrotetracycline (aTc) in a strain that contains TetR (Figure 4B), *BLM10* under the control of Tet operator sites (pTetO7.1-BLM10) is highly expressed (“*BLM10* o/e”) (Azizoglu et al., 2021). In the absence of aTc, TetR fully suppresses Blm10 expression (“*blm10*Δ,” Figure 4A, 4B). In strains with an endogenous copy of *BLM10* and no aTc introduced, only WT levels of Blm10 are produced (“WT,” Figure 4C, 4D). Using these systems, we assayed the effect of Blm10 levels on proteostasis.

**Figure 4.**
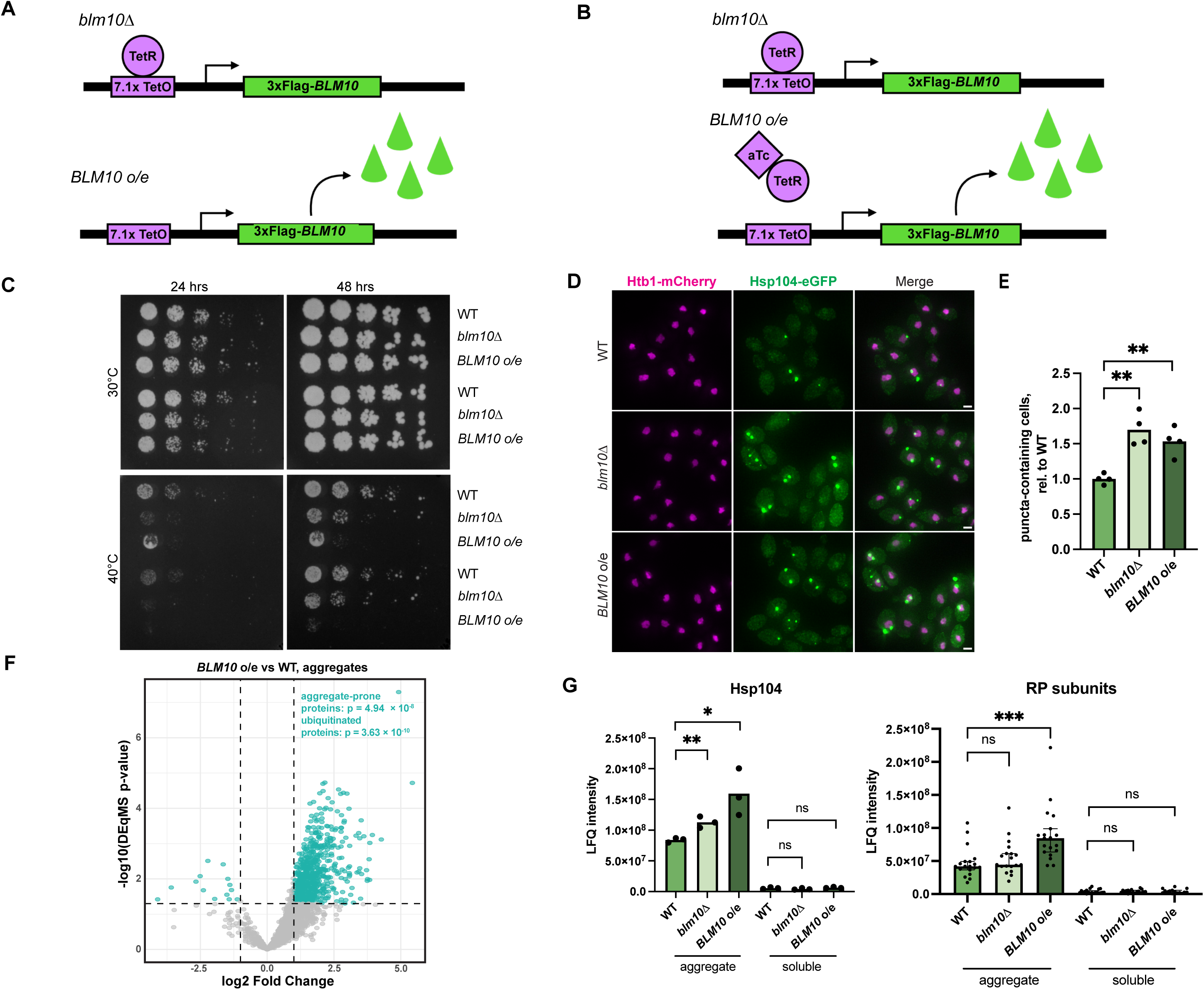
Either high or low Blm10 abundance disrupts proteostasis. (A,B) Schematic of aTc-inducible TetR/TetO expression systems ((Azizoglu et al., 2021) used in (C) and (D) respectively. (A) Expression of 3xFlag-*BLM10* is restricted when transcriptional repressor TetR is bound to TetO sites upstream of 3xFlag-*BLM10* (“*blm10*Δ”). Overexpression is achieved in the absence of TetR (“*BLM10* o/e,” Figure S4A). (B) Restriction of 3xFlag-*BLM10* occurs as in the same constructs as (A), “*blm10*Δ.” Overexpression is achieved when 5 μg/mL aTc is added, which releases TetR from binding TetO sites upstream of 3xFLAG-*BLM10* (“*BLM10* o/e,*”* Figure S4C). (C) Spot assay of WT, *blm10*Δ, and *BLM10* o/e cells on YPD plates grown at 30°C (top) and 40°C (bottom). Additional replicates in Figure S4B. (D) Fluorescence microscopy of Hsp104-eGFP marked aggregate foci in WT, *blm10*Δ, and *BLM10* o/e cells at log phase. Scale bar: 2 μm. (E) Ratio of puncta-containing cells relative to WT. 266 ≤ n ≤ 477 cells counted per genotype, 4 replicates. WT vs *blm10*Δ unpaired t test, p = 0.0013 (**). WT vs *BLM10* o/e unpaired t test, p = 0.0025 (**). *blm10*Δ vs *BLM10* o/e is ns. (F) Volcano plot depicting the log2 fold change of protein abundance, measured by mass spectrometry, in the aggregate fractions collected from *BLM10* o/e cells and WT cells. Significant proteins (log2 fold change >=1, -log(adjusted p value) <= 0.05) are highlighted in teal. These proteins that accumulate more in the *BLM10* aggregates are enriched for aggregate-prone proteins (Wallace et al., 2015) and ubiquitinated proteins (Tagwerker et al., 2006). (G) Mass spectrometry data for Hsp104 and RP subunits for WT, *blm10*Δ, and *BLM10* cells in the aggregate and soluble fractions. Hsp104: WT vs *blm10*Δ unpaired t test, p = 0.0071 (**). WT vs *BLM10* o/e unpaired t test, p = 0.0272 (*). RP subunits: WT vs *BLM10* o/e unpaired t test, p = 0.0152 (*).

Both *blm10Δ* and *BLM10* o/e cells grow slower compared to WT when heat-stressed, suggesting that an optimal level of Blm10 is important for mitotic growth under these conditions (Figure 4C, Figure S4B). Heat sensitivity is a condition in which specific misfolded and aggregation-prone proteins accumulate (Verghese et al., 2012; Wallace et al., 2015), hence we tested for markers of aggregation in *blm10Δ* and *BLM10* o/e cells. Indeed, we saw that Hsp104-eGFP-marked aggregate incidence was elevated by 1.5-fold in mitotic *blm10Δ* and *BLM10* o/e cells compared to WT controls (Figure 4D, 4E). Although the number of Hsp104 puncta was greater in *blm10Δ* and *BLM10* o/e cells, the overall abundance of Hsp104 protein in these strains remains unchanged (Figure S4C).

To investigate the nature of the aggregates formed in WT, *blm10*Δ, and *BLM10* o/e cells, we performed mass spectrometry on soluble- and aggregate-fractionated lysates (Figure S4D). These samples were collected in parallel and analyzed for Hsp104-marked aggregate puncta (Figure 4D). Surprisingly, few protein-level differences between WT and *blm10*Δ cells could be observed in the soluble and aggregate fractions (Figure S4D, S4E), but we observed a large cluster of 869 proteins that had increased abundance in the aggregate fraction of *BLM10* o/e cells compared to WT and *blm10*Δ cells (Figure S4D, green box).

Differential analysis of the aggregates revealed a large proportion of proteins significantly enriched in *BLM10* o/e cells compared WT (Figure 4F). Interestingly, hypergeometric enrichment analysis showed that aggregate-prone proteins, determined by their loss of solubility when cells are heat shocked, and proteins identified in a proteome-wide ubiquitin-profiling dataset (Tagwerker et al., 2006) were enriched in this group (Figure 4F), suggesting that these, among other proteins (Figure S4F), are constituents of aggregates specifically found in cells overexpressing *BLM10*. This is consistent with the observation that high Blm10 interferes with degradation of 26S target proteins (Figure 3C, 3D). We also observed that Hsp104 was specifically enriched in the aggregate-fractionated samples of *blm10*Δ and *BLM10* o/e cells, concordant with our previous microscopy data (Figure 4G, 4D). Intriguingly, an increase in abundance for all RP proteins was also specifically seen in the aggregate fractions of the *BLM10* o/e cells (Figure 4G). This is consistent with the model that an excess of Blm10 can displace RPs from the CP for binding (Figure 2G).

### Blm10 safeguards gamete quality

As reported previously, *blm10Δ* cells are overall healthy, as assessed by their similar mitotic rates compared to WT cells (Figure S5A, (Schmidt et al., 2005)). *blm10Δ* cells also progress through meiosis at a normal rate, sporulate as efficiently as WT cells, and a majority of the spores formed are viable (Figure S5B, S5C, S5D, (Iwanczyk et al., 2006)). The *blm10Δ* cells that we generated in the SK1 yeast background were respiration-competent, although this does not seem to be true of these mutants in the reference BY4741 strain background (Figure S5D, (Tar et al., 2014)). These data suggest that Blm10 is dispensable during mitotic growth in rich media, and for the completion of meiosis.

Since Blm10 appears to be dispensable for meiotic progression and spore formation, has roles in maintaining proteostasis, and is upregulated towards the end of gametogenesis, we hypothesized that the function of Blm10 in this context is to promote gamete fitness. Indeed, we observed that in a spore growth assay, in which individual spores are separated by microdissection and grown on rich media plates, *blm10*Δ spores produce colonies that are smaller than WT, indicating that they have a mild growth defect (Figure 5A). Consistently, time-lapse imaging of spore germination revealed that *blm10*Δ cells take longer to resume mitotic growth than WT controls, as assessed by timing with which the first bud is seen (Figure 5B).

**Figure 5.**
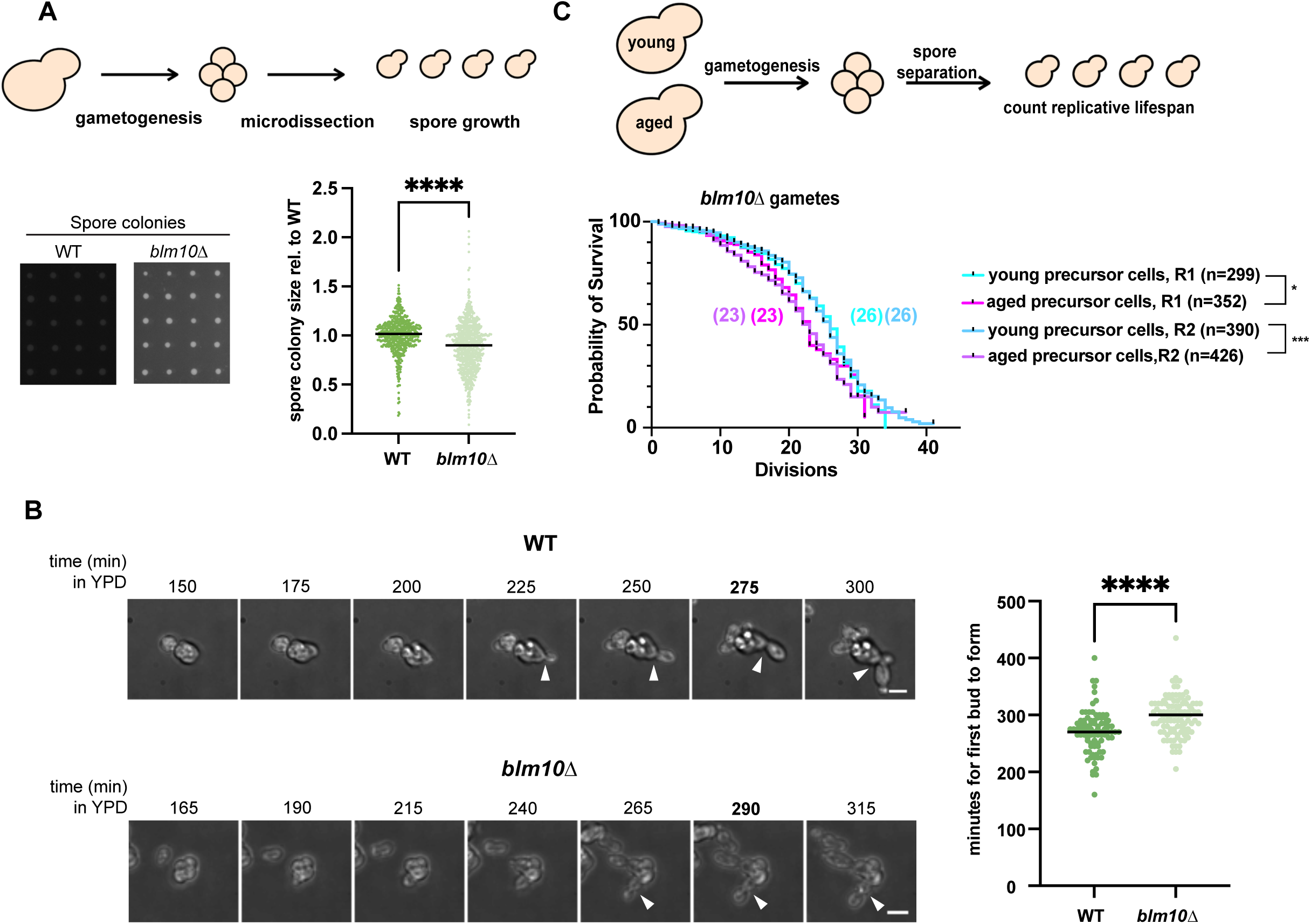
Blm10 safeguards gamete quality. (A) Schematic of spore germination assay. WT and *blm10*Δ cells undergo gametogenesis, and the resulting spores (gametes) are microdissected on YPD plates. Spores are allowed to grow at 30°C, then colony size is measured after 48 hours. Quantification of spore size for WT and *blm10*Δ cells relative to the average size of WT spores, with line at the median, is shown. WT: n = 537; median = 1, *blm10*Δ: n = 695; median = 0.8867. Welch’s t test, p < 0.0001. (B) Timelapse imaging of spore germination in liquid YPD for WT and *blm10Δ* cells. Buds form from spores when grown in rich media (white arrows, scale bar: 6 μm). Quantification for the time it takes for the first bud to form from a spore is shown (right, with line at median). WT median = 270 minutes, *blm10*Δ median = 300 minutes. WT: n = 86, *blm10*Δ: n = 107. Unpaired t test, p < 0.0001. The bolded timepoint in the representative images (left; WT = 275 minutes, blm10Δ = 290 minutes) is used for scoring. (C) Schematic for meiotic rejuvenation assay, as described in (Spiri et al., 2025). Cells are sorted into young and aged populations, undergo gametogenesis, and the resulting spores are separated. Replicative lifespan (the number of daughter cells that a mother cell can produce) is counted from timelapse imaging of spores trapped in microfluidic chips. Survival curves of *blm10*Δ cells from young and aged precursor cells are shown below. Median lifespan for replicate 1 and 2 (R1 and R2) gametes from young *blm10Δ* cells is 26. Median lifespan for replicate 1 and 2 (R1 and R2) gametes from aged *blm10Δ* cells is 23. R1 *blm10*Δ young precursor cells vs aged precursor cells p = 0.0272 (*), measured by log-rank Mantel-Cox test. *blm10*Δ young precursor cells vs aged precursor cells p = 0.0003 (***), measured by log-rank Mantel-Cox test. Survival curves of control gametes and the replicative lifespan of sorted young and aged populations, before and after gametogenesis, are found in Figure S6A and S6B.

Aging is a natural context in which the proteome is challenged, as aged cells accumulate damage, including protein aggregates (Klaips et al., 2017; Labbadia & Morimoto, 2015). Gametogenesis has the remarkable property of removing age-associated cellular damage, which manifests in the resetting of replicative lifespan for all four gametes produced from an aged precursor cell (Ünal et al., 2011). This involves quality control mechanisms that rejuvenate cellular contents so that damaged content in aged precursor cells does not get inherited into the gametes, such that they can start life “young” (King et al., 2019; Sing et al., 2022; Xiao & Ünal, 2025; Zaffagnini et al., 2024). We recently developed a robust and sensitive platform to assay gametogenic rejuvenation. In short, precursor cells are aged, undergo gametogenesis, and the resulting spores are separated and analyzed in microfluidic traps to assess their replicative lifespan (Spiri et al., 2025). Excitingly, gametes derived from aged *blm10*Δ precursor cells have a reproducibly significantly shorter replicative lifespan compared to young *blm10*Δ precursor cells. This is in contrast to what is seen in wild-type cells, in which both young and aged precursor cells produce gametes with equivalent replicative lifespan (Figure S6A, (Ünal et al., 2011)), indicating that Blm10 is needed for full lifespan resetting of aged precursor cells relative to young during gametogenesis (Figure 5C). Given Blm10’s role as a proteasome activator and the accumulation of protein aggregates in its absence, we conclude that Blm10 promotes the health and integrity of gametes by safeguarding proteostasis, perhaps by targeting misfolded, aggregation-prone proteins for degradation.

### Blm10-proteasomes physically interact with aggregation-prone proteins

To identify Blm10 interactors that may point to specific molecular functions of Blm10-proteasomes, we first immunoprecipitated FLAG-tagged Blm10-proteasomes. As we are interested in interactors that are unique to Blm10-proteasomes, rather than interactors that may be interacting with the RP in Blm10-CP-RP complexes, we performed a secondary IP on 6HIS-Rpn11 to differentiate between the “RP-containing” and “Blm10-only” complexes, which we then subjected to mass spectrometry analysis (Figure 6A). These purifications were performed in the presence of proteasome inhibitors MG132 and carfilzomib (CFZ) to stabilize potential substrates associated with the proteasome. Reproducibility was confirmed by replicate analysis (Figure S7A, S7B). We also observed that the amount of Blm10 in each sample was similar, and that Rpn11 is more abundant in the RP-containing proteasomes compared to the Blm10-only proteasomes (Figure S7C). As expected, we observed cyclin Cln2, a known proteasome target, to be enriched in the RP-containing complexes compared to the Blm10-only complexes (Figure 6B). Analysis of proteins in the Blm10-only complexes revealed that aggregation-prone proteins, as determined by their loss of solubility when cells are exposed to heat, were enriched ((Wallace et al., 2015), Figure 6B). GO term analysis revealed that proteins enriched in the Blm10-only complexes are known to be involved in protein folding, translation, and RNA binding, which are known to be enriched for intrinsically disordered regions and tend to be aggregate-prone (E. Gomes & Shorter, 2019), Figure 6B, C)).

**Figure 6.**
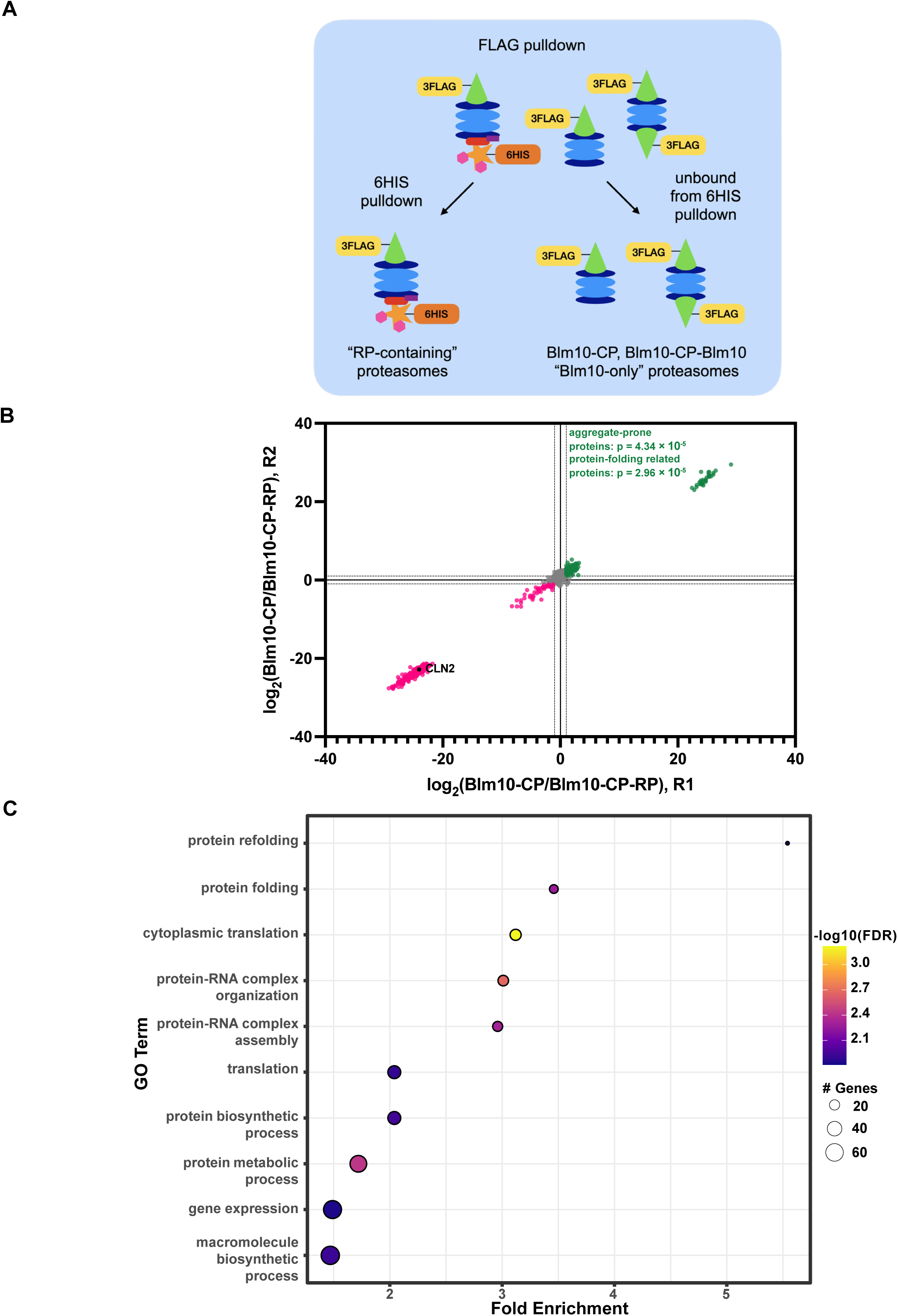
Blm10-proteasomes associate with aggregate-prone proteins and protein-folding factors. (A) Schematic of pulldown experiments to isolate “RP-containing” proteasomes (left) and “Blm10-only” proteasomes (right). 3xFLAG-Blm10 proteasomes were first isolated from mitotically growing cells treated with 50 μM MG132 and 20 μM carfilzomib to preserve substrate interactions, followed by a secondary pulldown on 6His-Rpn11 to separate Blm10-CP-RP proteasomes from the rest of the Blm10-proteasome pool, leaving Blm10-CP and Blm10-CP-Blm10 proteasomes in the unbound fraction. RP-containing proteasomes and Blm10-only proteasomes were then analyzed by mass spectrometry. (B) Replicate scatter plot of log2 fold changes of proteins measured in the Blm10-only sample over the RP-containing sample (R1: replicate 1, R2: replicate 2). Green: proteins enriched in Blm10-proteasomes over RP-containing proteasomes (120). Pink: proteins enriched in RP-containing proteasomes over Blm10-containing proteasomes (303). Aggregate-prone proteins ((Wallace et al., 2015) p = 4.34 × 10^-5^) and protein-refolding proteins (p = 2.96 × 10^-5^) are enriched in Blm10-only proteasomes over RP-containing proteasomes. (C) Gene ontology (GO) analysis for biological processed enriched in Blm10-only proteasomes over RP-containing proteasomes.

### Blm10 interacts in multiple stable configurations with the CP

To investigate potential cofactors of Blm10-proteasomes, we solved cryo-EM structures of purified Blm10-proteasomes from mitotic cells. To enrich for Blm10-proteasomes, cells were induced to overexpress 3xFlag-Blm10 by use of an anhydrotetracycline (aTc)-inducible TetR/TetO system (Azizoglu et al., 2021). Blm10 has been previously characterized to sit atop the core proteasome, and we also observe this structure, which we term the “rigid” cap structure (Guan et al., 2020; Iwanczyk et al., 2006; Kaur et al., 2024; Qian et al., 2013; Sadre-Bazzaz et al., 2010; Schmidt et al., 2005; Toste Rêgo & da Fonseca, 2019). However, we also observed a structure in which Blm10 is rotated counterclockwise and upwards while bound to the CP, as if on a hinge, so that the inside of the Blm10 dome is solvent facing (Figure 7A). We term this the “swivel cap.” A substantial proportion (25%) of Blm10-proteasomes showed Blm10 positioned in this swiveled conformation.

**Figure 7:**
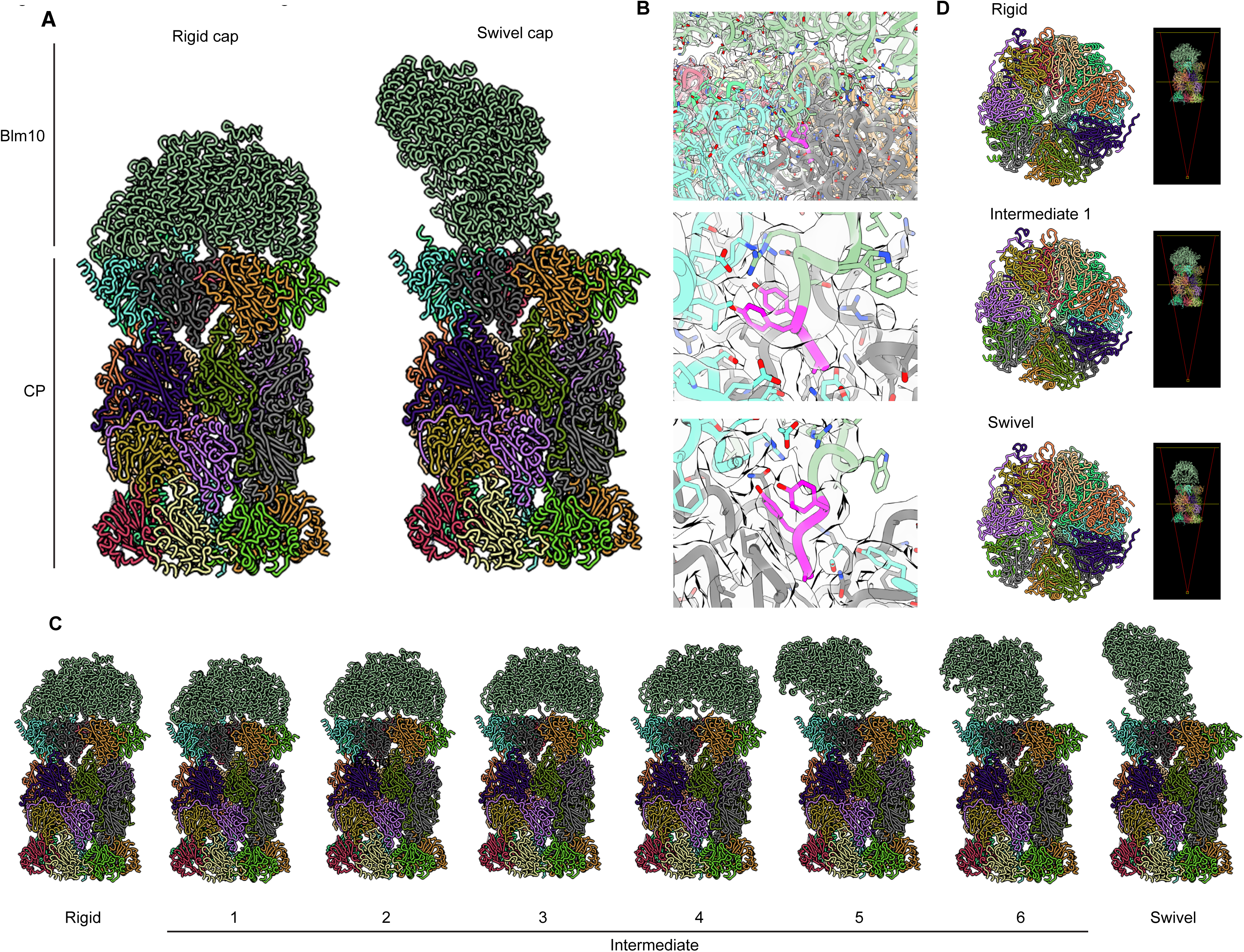
Blm10 can be seen in “rigid” and “swivel” conformations on the CP. (A) Models, determined from cryo-EM, of the “rigid” (left) and “swivel” (right) conformations of Blm10 atop the CP. Models are built for the top Blm10 and complete 20S CP. (B) Views of Blm10 YYA (C-terminal HbYX motif) bound in the inter-subunit pocket between α5 (cyan) and α6 (black) when Blm10 is in the swiveled conformation, in the same manner that YYA is docked into the α-ring when Blm10 is in the rigid conformation. (C) 6 intermediates resolved between the Blm10 rigid and swiveled state. (D) Gate status of the rigid cap, firstintermediate, and swivel cap, viewed through the internal degradation chamber of the CP (representations to the right). Note that the α-ring is opened in the rigid conformation and closed in the first intermediate and in the swiveled state.

Notably, the C-terminal HbYX motif of Blm10, which allows for binding to the CP and opening of the α-ring, remains docked in a similar position whether the configuration is swiveled or not (Figure 7B), suggesting that the mode of binding remains the same. We also resolved 6 intermediate states between the rigid and swivel cap (Figure 7C). It appears that parts of the repeating solenoids in Blm10’s structure make contacts with the CP when swiveled (Figure 7A, S9C). While it is known that the α-ring is open when the rigid cap is bound, interestingly, in the swivel state, the α-ring is closed (Figure 7D, “Rigid,” “Swivel”). Furthermore, the α-ring appears closed in all swivel conformations (Figure 7D, “Intermediate 1”). These data reveal that Blm10 can bind to the CP in alternative manner, and suggest that a series of controlled switches occur from the rigid to swivel state, taking Blm10 proteasomes between a rigid cap configuration with an open α-ring and a configuration in which the Blm10 cap is swiveled open but the CP α-ring is closed.

### 14-3-3 adaptor proteins Bmh1/2 stably binds to Blm10

For the rigid Blm10-proteasome complexes, we identified a complex with a hook-like structure bound to the top of the Blm10 dome. This was observed in about 40% of the rigid Blm10-proteasome structures and was also present in all other configurations including the swivel form (Figure 8A). To determine the identity of the hook-like structure, a portion of the purified sample prepared for cryo-EM was analyzed by mass spectrometry. Excitingly, 14-3-3 proteins Bmh1 and Bmh2 were identified amongst the top hits in these samples, which were otherwise composed of primarily proteasome subunits (Figure 8B). We could not differentiate between the Bmh1 and Bmh2 isoforms from the EM density, indicating that a mixture of both cofactors may be bound. However, we found that available structures of Bmh1 docked well inside initial density maps, and atomic modeling assigned the hook-like structure to Bmh1 (Figure 8A). The region of Bmh1 that binds to Blm10 (residues 155-233) is also identical to that of Bmh2, suggesting that both isoforms can stably interact with Blm10-proteasomes.

**Figure 8.**
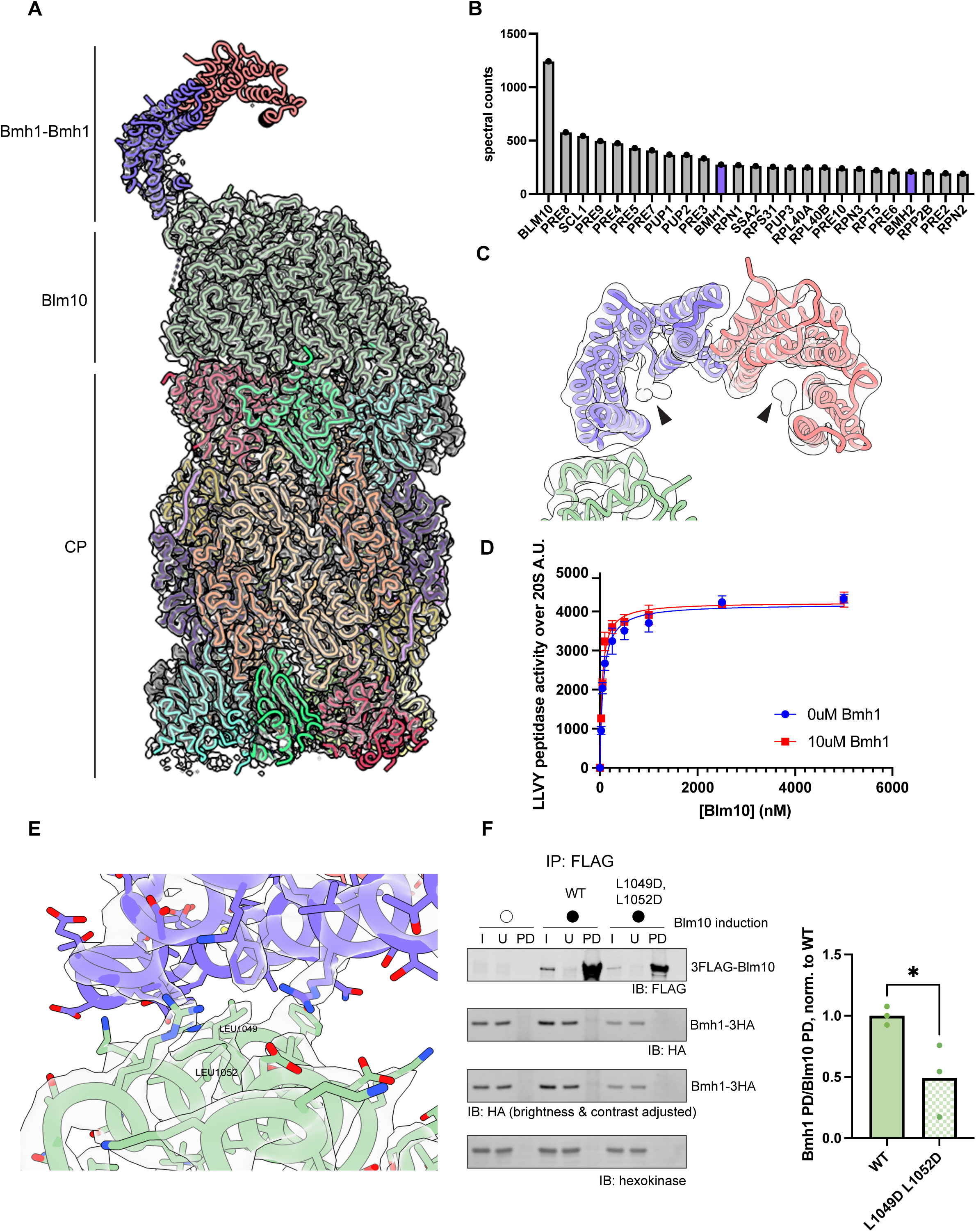
Cryo-EM structure of 14-3-3 protein Bmh1 bound to the Blm10-proteasome. (A) Structure of Bmh1-Blm10-CP. Bmh1 homodimer is modeled. Model is built for the top Blm10 and Bmh1 and complete 20S CP. (B) Relative quantification of protein abundance for the top hits identified in purified 3FLAG-Blm10 proteasomes determined by label-free quantification mass spectrometry. Gray: proteasome components, purple: Bmh1, and paralog Bmh2. (C) Density in the phospho-binding pocket, indicated by black arrows, of Bmh1-Bmh1 homodimer (pink and purple) bound atop Blm10 (green). (D) Proteasome chymotryptic activity measured upon Blm10 titration to the 20S CP (same experiment as in Figure 2F, only 0 µM Bmh1 shown). Upon the addition of 10 µM Bmh1, no change in proteolytic activity is observed. (E) Blm10-Bmh1 binding interface, with Blm10 in green and Bmh1 in purple. Key residues L1049 and L1052 are labeled on Blm10. (F) Co-IP of 3FLAG-Blm10 in mitotic cells. Quantification of the amount of Bmh1-3HA that co-immunoprecipitated with 3FLAG-Blm10 is shown. Replicates shown in S8C. I = input, U = unbound, PD = pulldown.

Bmh1 is a member of the 14-3-3 family, which bind phosphorylated residues (Johnson et al., 2010; Obsil & Obsilova, 2011; Yaffe et al., 1997). 14-3-3 proteins can exist in hetero-or homodimeric complexes, forming a horseshoe shape, in which phosphopeptides are bound in the C-terminal “crook” of the horseshoe (Obsil & Obsilova, 2011; Yaffe et al., 1997). In yeast, Bmh1 and Bmh2 are the sole 14-3-3 isoforms (Obsil & Obsilova, 2011) and Bmh1 is likely the more dominant isoform as assessed by proteomic measurements (Ho et al., 2018; Kumar, 2018). 14-3-3 proteins can function as a scaffold that facilitates protein-protein interactions, with crystal structures demonstrating that 14-3-3 proteins can accommodate two substrates in each “crook” of its horseshoe-like structure (Obsil et al., 2001; Ottmann et al., 2007). Recently, a holdase-type chaperone role has been proposed for 14-3-3 proteins, as their interaction with disordered clients, including RNA-binding proteins, prevents client aggregation (Segal et al., 2023).

Excitingly, we were able to observe additional density inside the Bmh1 phospho-pocket in the context of Bmh1-Blm10-proteasome complexes (Figure 8C, black arrows). While we were not able to further resolve the density, we hypothesize that it could represent a bona fide Bmh1-phosphosubstrate. Therefore, we investigated if known Bmh1 interactors co-purified with Blm10-proteasomes. Analysis of proteins enriched in the purified mitotic Blm10-proteasome samples we previously collected (Figure 6) revealed significant overrepresentation of established Bmh1 interactors (4.29E-3 p-value by hypergeometric enrichment analysis; Saccharomyces Genome Database). These data suggest that Bmh1 can bind with known interactors and Blm10-proteasomes at the same time and may suggest that this is how substrates are recruited to Blm10 proteasomes.

We found that the CP α-ring on the Bmh1-Blm10-proteasome is open to a similar degree with and without Bmh1 (Figure S8A). Consistently, addition of Bmh1 does not change the proteolytic activity of Blm10-proteasomes (Figure 8D). Noticeably, a loop on Blm10 that projects inside the dome undergoes a distinct conformational change dependent on Bmh1 binding (Figure S8B, Video 1), however the significance of this change requires further investigation.

### Two central leucines maintain the interaction between Blm10 and Bmh1/2

Analysis of Bmh1-Blm10-proteasome structures allowed us to define the binding interface between Blm10 and Bmh1 to include Blm10 residues 174-180 and 1045-1056, representing two helices that are located by the top of the Blm10 dome, which interact with Bmh1 residues 153-233 (Figure 8E). This region is distinct from the phospho-peptide binding region of Bmh1, and surprisingly, it appears to be mediated by primarily hydrophobic interactions. We hypothesized that residues L1049, and L1052 were key for maintaining this interaction, as they are the residues with the most direct contact to Bmh1 in the structures (Figure 8E). L1049 and L1052 are located on one of the Blm10 helices that participates in the interaction. We generated point mutants for these residues (“L1049D, L1052D”), to investigate if mutations to aspartate would disrupt the hydrophobic interactions of the interface and diminish binding of Bmh1 to Blm10. Co-IP of the Blm10 L1049D, L1052D mutant revealed diminished binding to Bmh1 (Figure 8F). While we did not observe complete abolishment of binding, we hypothesize that additional residues in the interface, including perhaps the involvement of other Blm10 segments that are unable to be resolved structurally, may also contribute to binding.

## Discussion

Here, we report that Blm10 is an upregulated proteasome activator during gametogenesis that functions to regulate proteostasis and ensure gamete quality. Blm10-proteasomes, including Blm10-CP-Blm10, Blm10-CP, and hybrid Blm10-CP-RP complexes are populated during gametogenesis. Blm10 is a weak proteasome activator, which binds to the CP in a concentration-dependent manner and is able to compete with the RP for binding. Either too little or too much Blm10 lead to proteostatic defects. Blm10 is critical in supporting normal gamete fitness and rejuvenation. Structural data show that Blm10 binds in novel configurations to the CP and that it stably associates with 14-3-3-proteins, which may define the clients of this poorly understood proteasome type.

We propose a model by which an optimal amount of Blm10 is important for maintenance of proteostasis, based on a balance of its effects on the abundance of 26S species and its influence on the fate of a subset of poorly folded and/or aggregation-prone substrates (Figure 9). Thus, in the absence of Blm10, a subset of misfolded proteins is not properly targeted for degradation. This could explain the heat sensitivity of *blm10*Δ cells, their accumulation of Hsp104 foci, and their inability of these cells to rejuvenate during gametogenesis despite minimal changes in the bulk levels of individual proteins (Figure 9). Consistently, analysis of proteins associated with Blm10-proteasomes showed enrichment in aggregate-prone proteins and chaperones (Figure 6B, 6C). A previous study has shown that toxic Huntingtin aggregates (Htt103Q) accumulate in *blm10Δ* yeast cells and are degraded by Blm10-proteasomes *in vitro*, suggesting a role for Blm10-proteasomes in the quality control of aggregation-prone proteins (Aladdin et al., 2020). Further study is necessary to definitively link putative substrates of Blm10-proteasomes to proteostatic defects.

**Figure 9.**
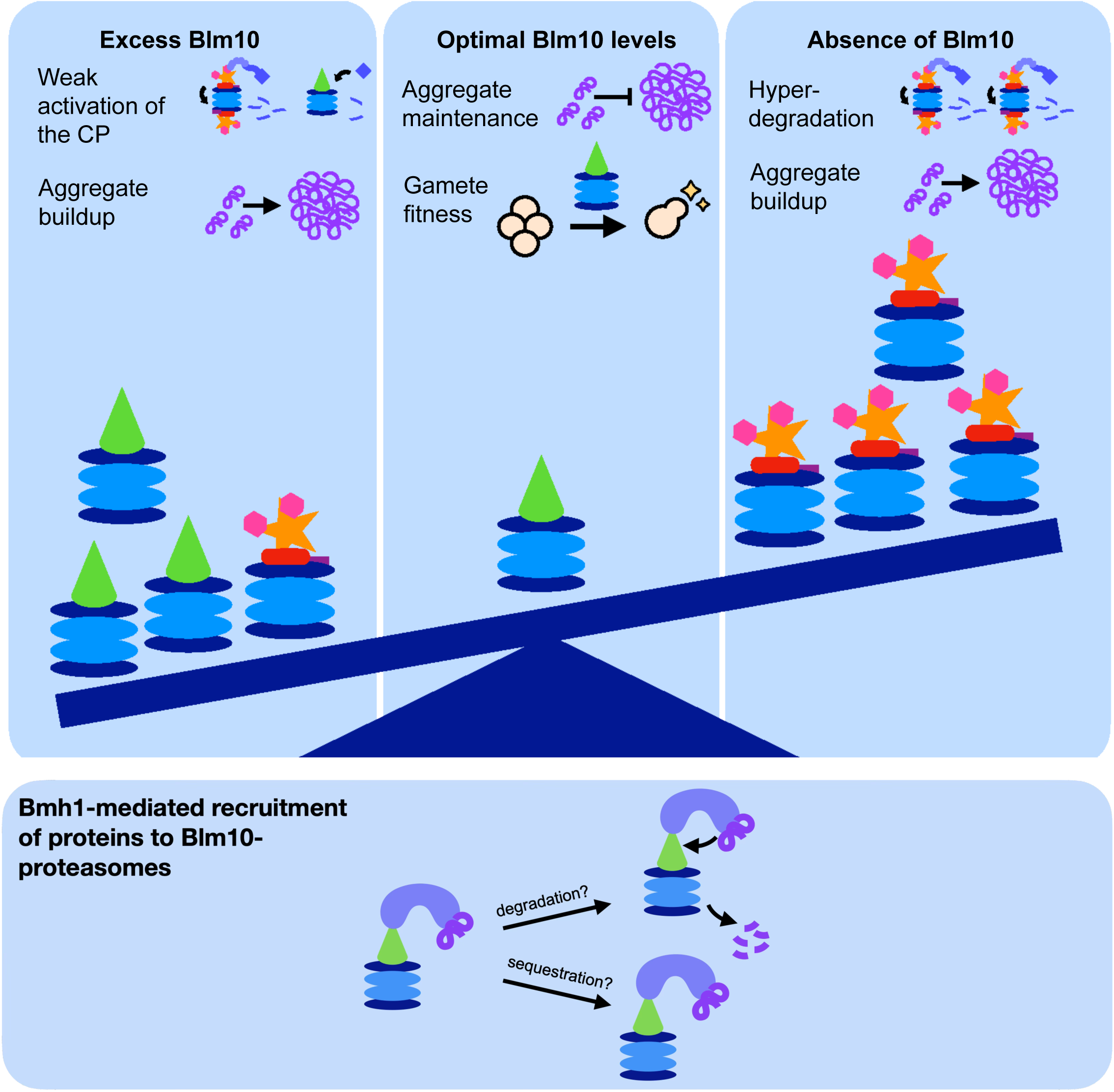
Model for Blm10-proteasome function. (A) Model for the role of Blm10-proteasomes in proteostatic regulation and gamete fitness. In conditions of excess Blm10, bound CP’s have weak proteolytic activity, and protein aggregates accumulate. Additionally, bound RP can be displaced from the CP. In conditions in which Blm10 is absent, 26S proteasomes hyperdegrade certain substrates, and aggregates that may be targeted by Blm10 accumulate. In conditions of optimal Blm0 abundance, aggregates are prevented, and gamete fitness is maintained. Lastly, 14-3-3 protein Bmh1 binds to Blm10-proteasomes. Bmh1 may recruit substrates for either degradation by the complex, or for sequestration without a degradative outcome.

When there is an excess of Blm10, the level of 26S proteasome complexes may decrease, as Blm10 is capable of displacing the RP. This could explain the buildup of a subset of misfolded proteins, evidenced by increased Hsp104 foci and heat sensitivity of such cells that we observe with Blm10 overexpression (Figure 9). In vivo competition between Blm10 and the RP for CP binding would also be consistent with the hyper-degradation of Sic1 and other 26S proteasome targets in cells lacking Blm10. Strikingly, proteins identified as targets of ubiquitination (Tagwerker et al., 2006) were enriched in aggregates of cells overexpressing *BLM10* (Figure 4F, 4G), also consistent with a model in which Blm10 displaces the RP for binding to the CP. Our study and others have shown that a diversity of proteasome complexes with different catalytic functions are found within cells and can vary between tissues (Dahlmann et al., 2000; Fabre et al., 2014; A. V. Gomes et al., 2009; Sahoo et al., 2024). It is an attractive hypothesis that optimal stoichiometry of different proteasome complexes ensures that the homeostatic needs of the cell or organism are met. Consistently, upregulation of the immunoproteasome correlates with improved survival in breast cancer, and increased sensitivity to proteasome inhibition in acute myeloid leukemia M5 cells (Rouette et al., 2016).

The structures that we report here may be the first step toward understanding two previously confounding features of Blm10 proteasomes: how substrates could enter the CP when Blm10 is bound and how Blm10-proteasome substrates are determined. We resolved cryo-EM structures of a new class of Blm10-proteasomes, with Blm10 partly swiveled away from the CP (Figure 7). We speculate that substrates could become trapped underneath the swiveled Blm10 dome and could access the degradative chamber of the CP when Blm10 swivels shut to the rigid state, and the α-ring gate opens. Since the gate remains closed with swiveled Blm10, this suggests that the rigid to swivel transition is a controlled switch that regulates access to the CP. This could be advantageous for any non-target proteins in the vicinity, since a closed, exposed α-ring prevents premature degradation. Whether evidence for swiveled Blm10 proteasomes can be seen within cells is an exciting future area of investigation. Through proteomic and structural analysis, we also discovered 14-3-3 protein Bmh1 as a novel cofactor for Blm10-proteasomes. We found known Bmh1-interacting proteins to be enriched in purified Blm10-proteasomes, suggesting that Bmh1 acts as a scaffold that may recruit proteins to the Blm10-proteasome. 14-3-3 proteins have recently been shown to function as “holdase”-type chaperones, binding disordered proteins to prevent their aggregation (Segal et al., 2023), which is consistent with our observation that aggregate-prone proteins significantly co-purified with Blm10-proteasomes (Figure 6).

Why are Blm10-proteasomes enriched in late stages of gametogenesis? It is interesting to note that Blm10 has a mammalian ortholog, PA200 (Ustrell et al., 2002), which is upregulated during, and required for, gametogenesis in male mice (Khor et al., 2006; Sato et al., 2023). It is of great interest whether PA200 has a similar role as Blm10 in the maintenance of proteostasis in higher eukaryotes during gametogenesis. This is known to be a context in which clearance of aggregate proteins is of particular importance, as gamete quality determines the fate of future generations. Aging is an especially well-studied context in which proteostasis is overtaxed and protein aggregates accumulate (Hipp et al., 2019; Klaips et al., 2017; Labbadia & Morimoto, 2015). Gametogenic rejuvenation is known to clear these age-associated aggregates, at least in part through degradation, but the role of the proteasome in this clearance has been unclear (King et al., 2019; Sing et al., 2022; Ünal et al., 2011). It is also known that a substantial portion of the gamete proteome is degraded and resynthesized after completing gametogenesis (Eisenberg et al., 2018). One justification for the resetting of the gamete proteome is that it preserves gamete integrity, which in turn ensures an organism’s fitness over evolutionary time (Eisenberg et al., 2018). Blm10 is upregulated towards the end of gametogenesis and given the associations that we identified between Blm10 proteasomes and aggregation-prone proteins, it is tempting to speculate Blm10 safeguards the gamete proteome, a role that is particularly important in cells housing the type of damaged or aggregated proteins that accumulate in aged cells. Consistently, gametes lacking Blm10 display evidence of poor quality, and aged cells lacking the ability to express Blm10 in gametogenesis produce gametes with an incompletely reset replicative lifespan. Blm10 is one of the first factors known to be responsible for the remarkable rejuvenation that underlies gamete production, and understanding the specific substrates that are responsible for this function is an exciting future area of research.

## Acknowledgements

We thank James Olzmann, Nicholas Ingolia, Jeremy Thorner, and the entire Brar-Ünal Lab and Martin Lab for valuable discussions. We thank the UCSF CAT sequencing core and UC Berkeley QB3 mass spectrometry facility for assistance in mRNA-sequencing and mass spectrometry data collection. This work was supported by NIH R01 AG071869 to G.A.B. and M.J. E.D.-M. was supported by NIH/NINDS (K99NS135103). A.M. is supported by HHMI funding.

## Author contributions

M.C.R., D.R.R.O., G.A.B., M.J. and A.M. designed the research. M.C.R., D.R.R.O., J.A.C., S.S., E.D.-M., J.K.K., and Y.M. performed the experiments. M.C.R. and G.A.B. wrote the original draft of the manuscript. All authors edited and revised the final manuscript.

---

**Supplementary Figure 1.**
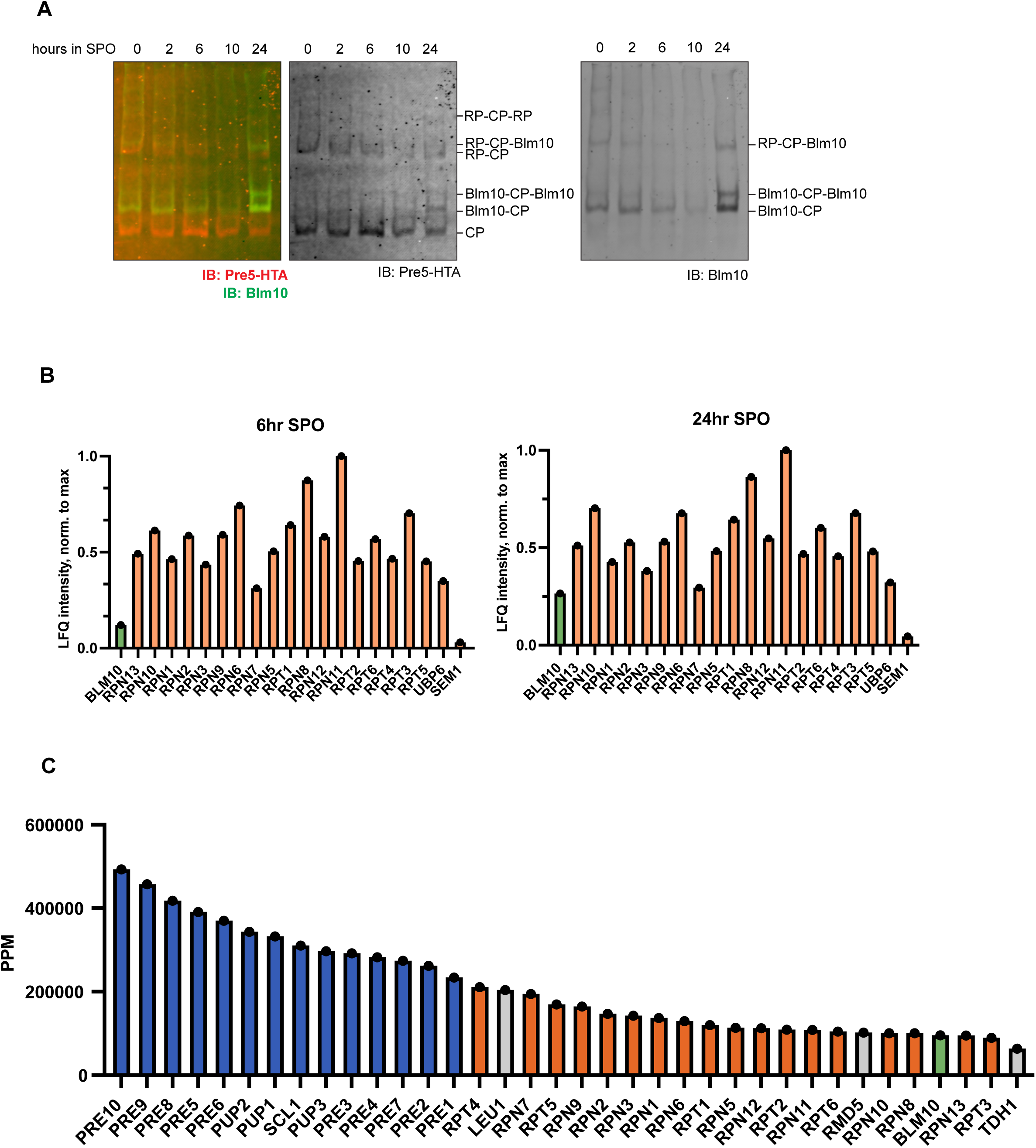
(A) Native immunoblot replicate for Pre5-HTA (CP, red) and Blm10 (green). (B) Mass spectrometry abundance of Blm10 (green) and all RP proteins detected (orange) at 6 hours and 24 hours, as in (Eisenberg et al., 2018). (C) Mass spectrometry abundance the top 50 proteins detected in purified Pre5-3FLAG proteasomes isolated from 24 hours in SPO. CP (blue), RP (orange), Blm10 (green), and other proteins (gray) are shown.

**Supplementary Figure 2.**
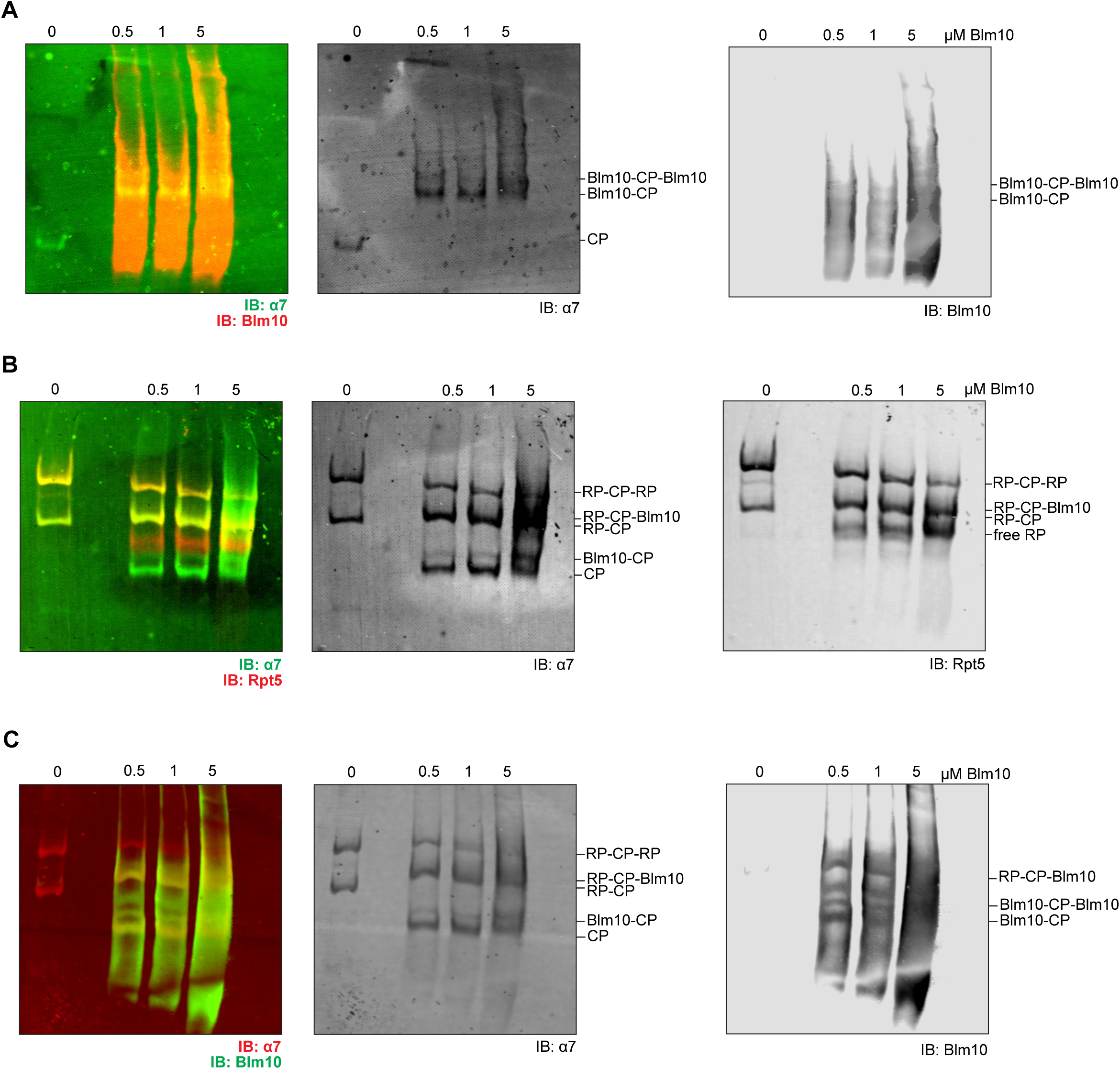
(A) Native immunoblot for Blm10 (red) and α7 (CP, green), for 25 nM purified CP + indicated [3x-Flag-Blm10] in Figure 2F. (B) Native immunoblot for Rpt5 (RP, red) and α7 (CP, green), for 100 nM purified 26S proteasome + indicated [3x-Flag-Blm10] in Figure 2G. (C) Native immunoblot for Blm10 (green) and α7 (CP, red), for 100 nM purified 26S proteasome + indicated [3x-Flag-Blm10] in Figure 2G.

**Supplementary Figure 3.**
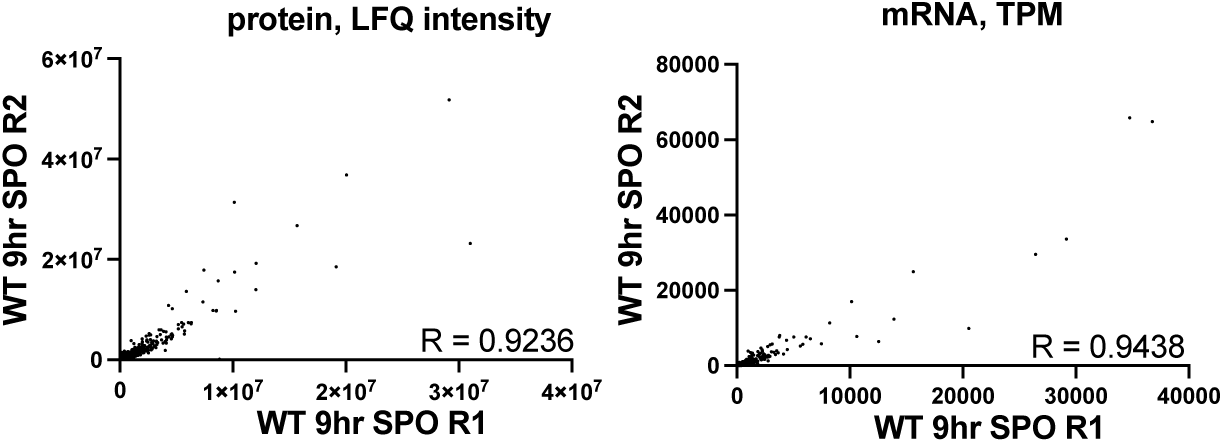
Scatterplots of replicate samples (R1, R2; 9 hours in SPO) for mass spectrometry data (left, label free quantification (LFQ) intensity) and mRNA-seq data (right, transcripts per million (TPM)).

**Supplementary Figure 4.**
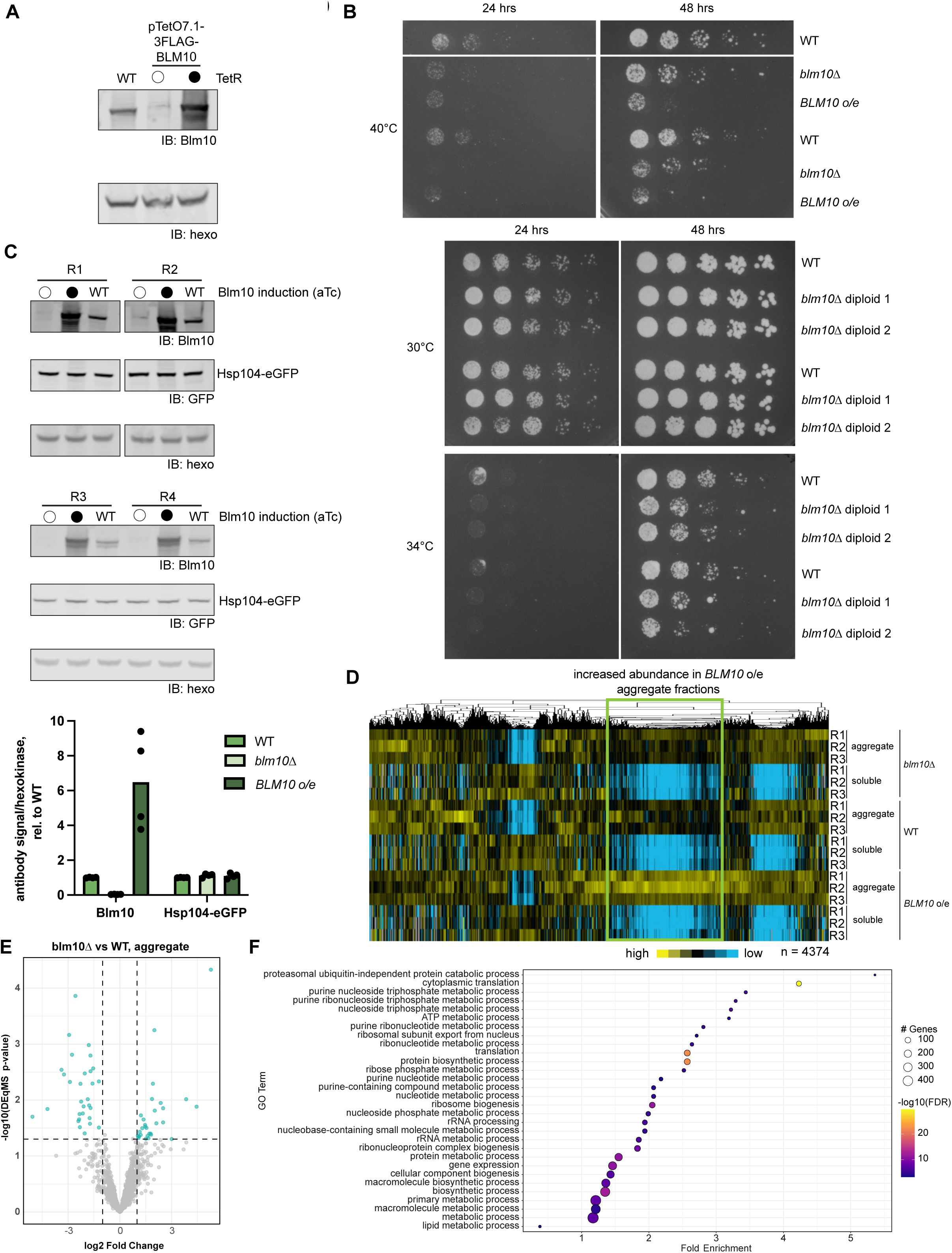
(A) Western blot of Blm10 produced from the absence of the TetR allele. (B) Replicates of spot assay shown in Figure 3B (top: replicate of the same strains used in Figure 4C. Bottom: replicates of diploid WT and blm10Δ cells). (C) Western blot of four replicates (R1-R4) analyzed in Figure 4D-F for Blm10 and Hsp104-eGFP levels. Quantification shown below. (D) Global representation of mass spectrometry (MS) data (label-free quantification intensity (LFQ), normalized across genes and hierarchical clustered) for matched fractionated samples from blm10Δ, WT, and BLM10 o/e cells analyzed in Figure 4D-G, S4C-F. 3 replicates (R1, R2, and R3) of aggregate and soluble fractions are shown for each genotype. Cluster containing proteins that exhibit increased abundance in BLM10 o/e aggregate fractions is highlighted within the green box. N = 4374 proteins quantified. (E) Volcano plot of diferentially abundant proteins in the aggregate fractions (n = 3) isolated for blm10Δ over WT. Teal dots represent proteins with p < 0.05. (F) Gene ontology (GO) analysis for biological processes for proteins enriched in the blm10Δ aggregate fraction over WT (shown in Figure 4F).

**Supplementary Figure 5.**
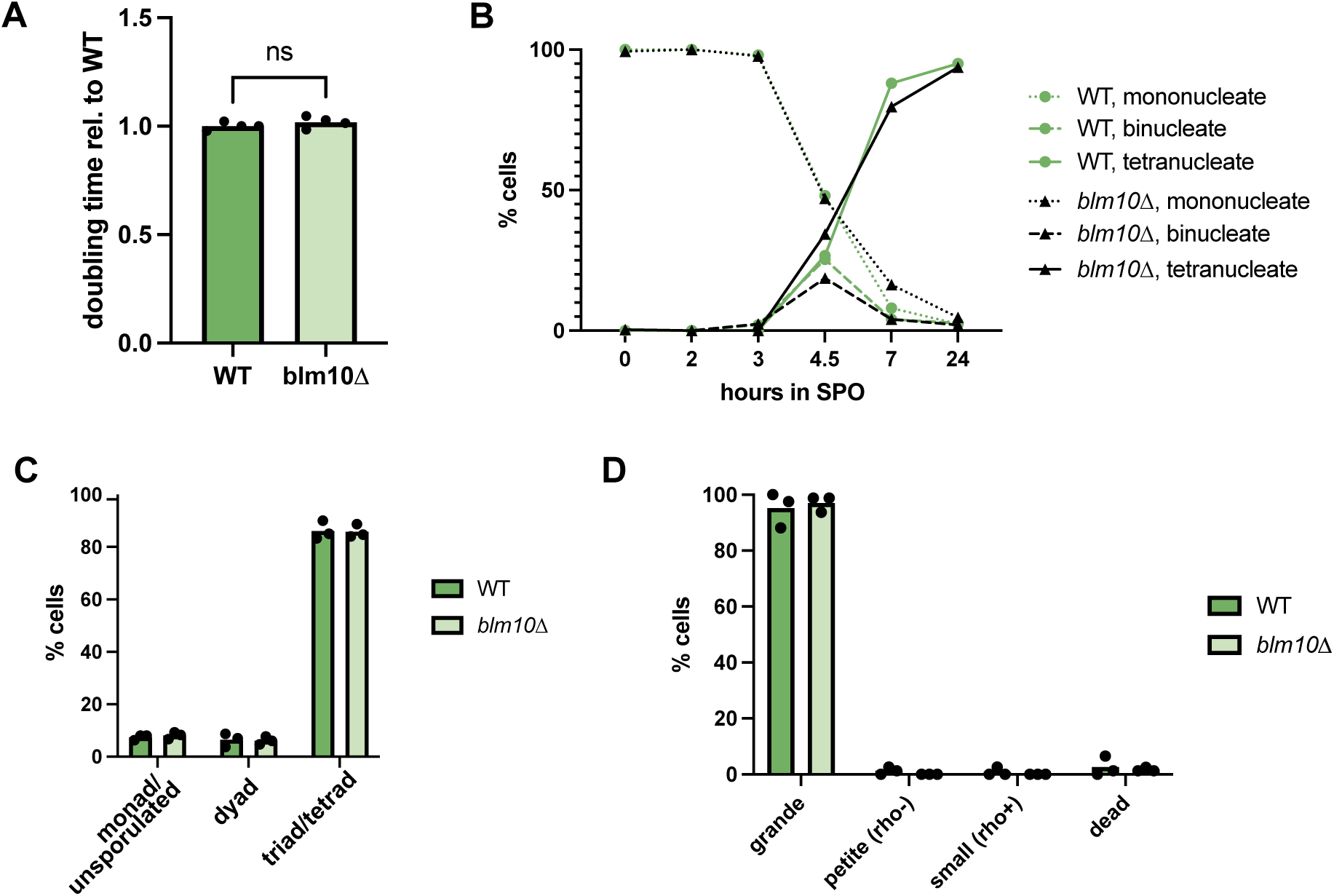
(A) Doubling time for WT and blm10Δ cells, relative to WT, in YPD at 30C. 4 replicates from growth curves shown. (B) Meiotic progression monitored by DAPI staining for WT (green) and blm10Δ (blue) cells. N = 300 cells per genotype counted per timepoint in SPO. (C) Sporulation eficiency for WT (green) and blm10Δ (blue) cells. N = 300 cells per genotype, 3 replicates. (D) Spore viability of WT, blm10Δ cells. Petite, rho-spores are classified as smaller colonies that cannot grow on replica plated YPG plates, which contain the non-fermentable carbon source glycerol. Small colonies that survive on YPG plates are rho+. WT: R1 n = 76, R2 n = 80, R3 n = 80; blm10Δ: R1 n = 80, R2 n = 80, R3 n = 80.

**Supplementary Figure 6.**
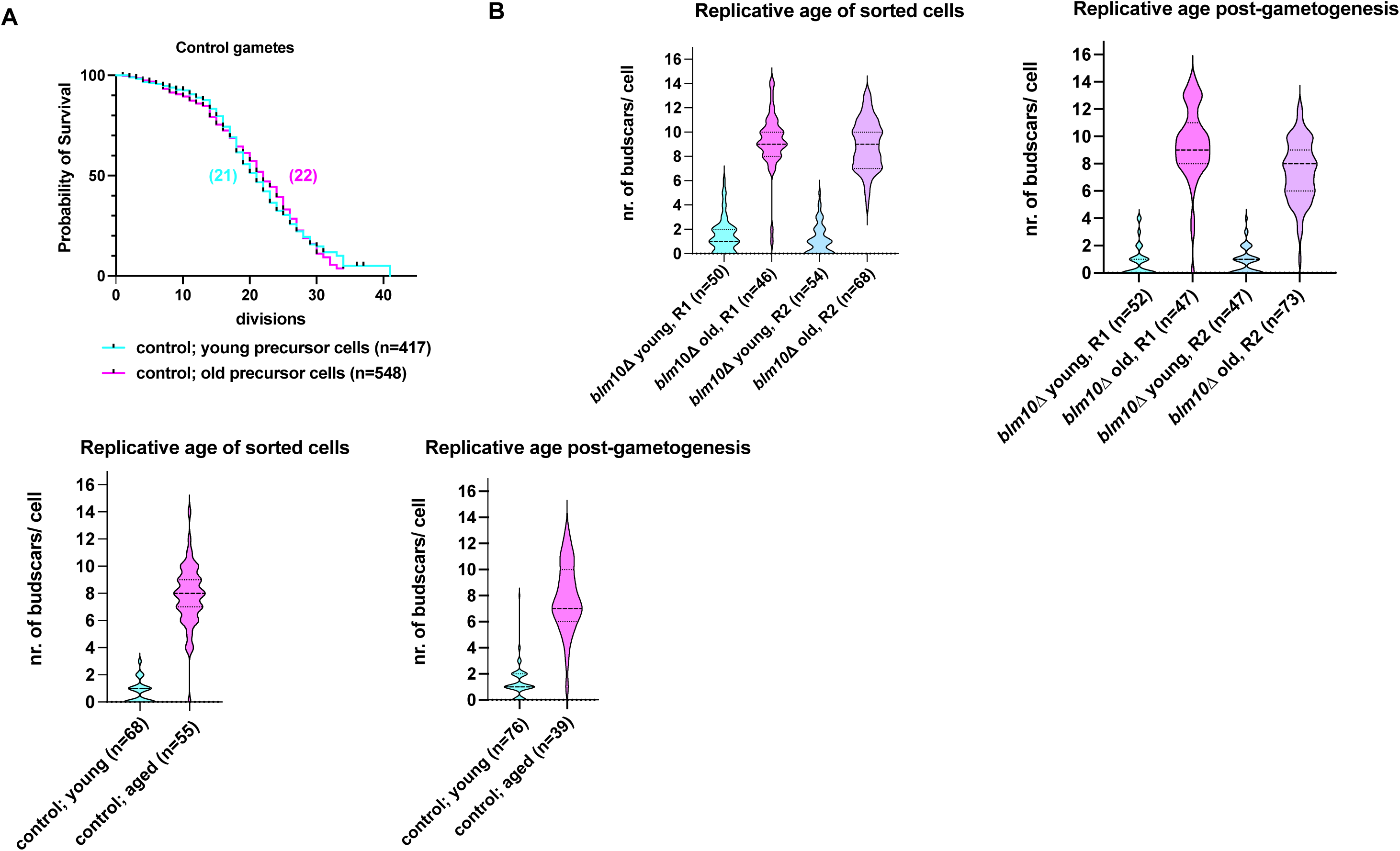
(A) Survival curves for control cells from the meiotic rejuvenation assay, as published in Spiri et. al 2025. Median lifespan for gametes from control young precursor cells is 21, median lifespan for gametes from control aged precursor cells is 22 (ns). Replicative age, measured by number of budscars per cell, of young and aged control gametes before gametogenesis (young: n = 68, aged: n = 55). Replicative age, measured by number of budscars per cell, of young and aged control gametes after gametogenesis (young: n = 76, aged: n = 39) (B) Replicative age, measured by number of budscars per cell, of young blm10Δ gametes before gametogenesis (R1: n = 50, R2: n = 54). Replicative age, measured by number of budscars per cell, of aged blm10Δ gametes before gametogenesis (R1: n = 46, R2: n = 68). Replicative age, measured by number of budscars per cell, of young blm10Δ gametes after gametogenesis (R1: n = 52, R2: n = 47). Replicative age, measured by number of budscars per cell, of aged blm10Δ gametes after gametogenesis (R1: n = 47, R2: n = 73).

**Supplementary Figure 7.**
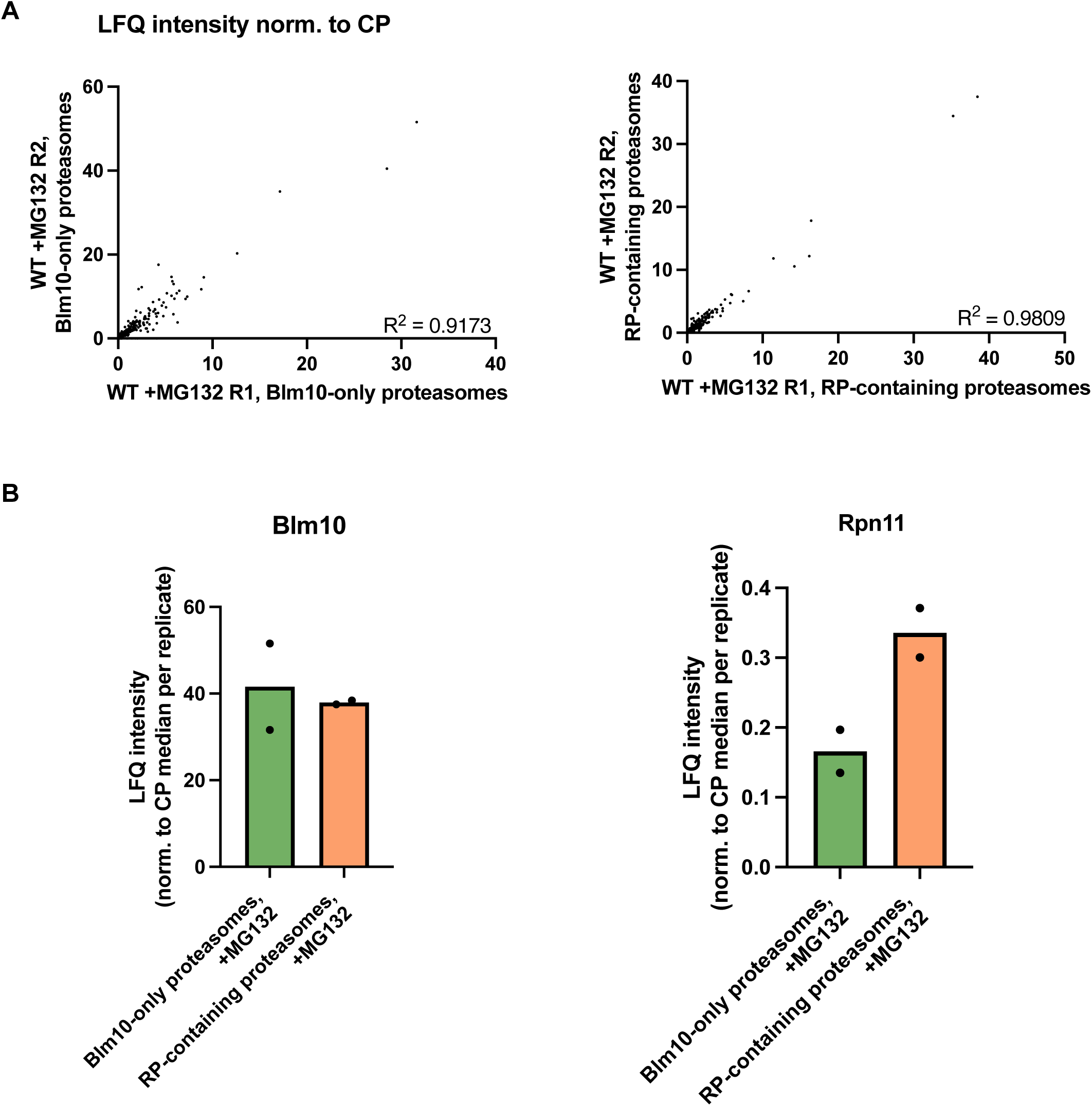
(A) Replicate analysis of LFQ mass spectrometry measurements of isolated Blm10-CP and Blm10-CP-Blm10 complexes (“Blm10-only proteasomes”) and isolated RP-CP-Blm10 proteasomes (“RP-containing proteasomes”) from WT cells treated with 50 μM MG132 and 20 μM carfilzomib (“+MG132”). Blm10-containing proteasomes: R2 = 0.9173, n = 2. RP-containing proteasomes: R2 = 0.9809, n = 2. (B) Protein abundance (LFQ intensity) of Blm10 and Rpn11 in Blm10-only proteasomes and RP-containing proteasomes, normalized to the median CP protein abundance measured per replicate. Abundance of Blm10 is comparable in both Blm10-only proteasomes and RP-containing proteasomes, whereas Rpn11 (which was N-terminally tagged with 6His to isolate RP-containing proteasomes) is enriched in RP-containing proteasomes.

**Supplementary Figure 8.**
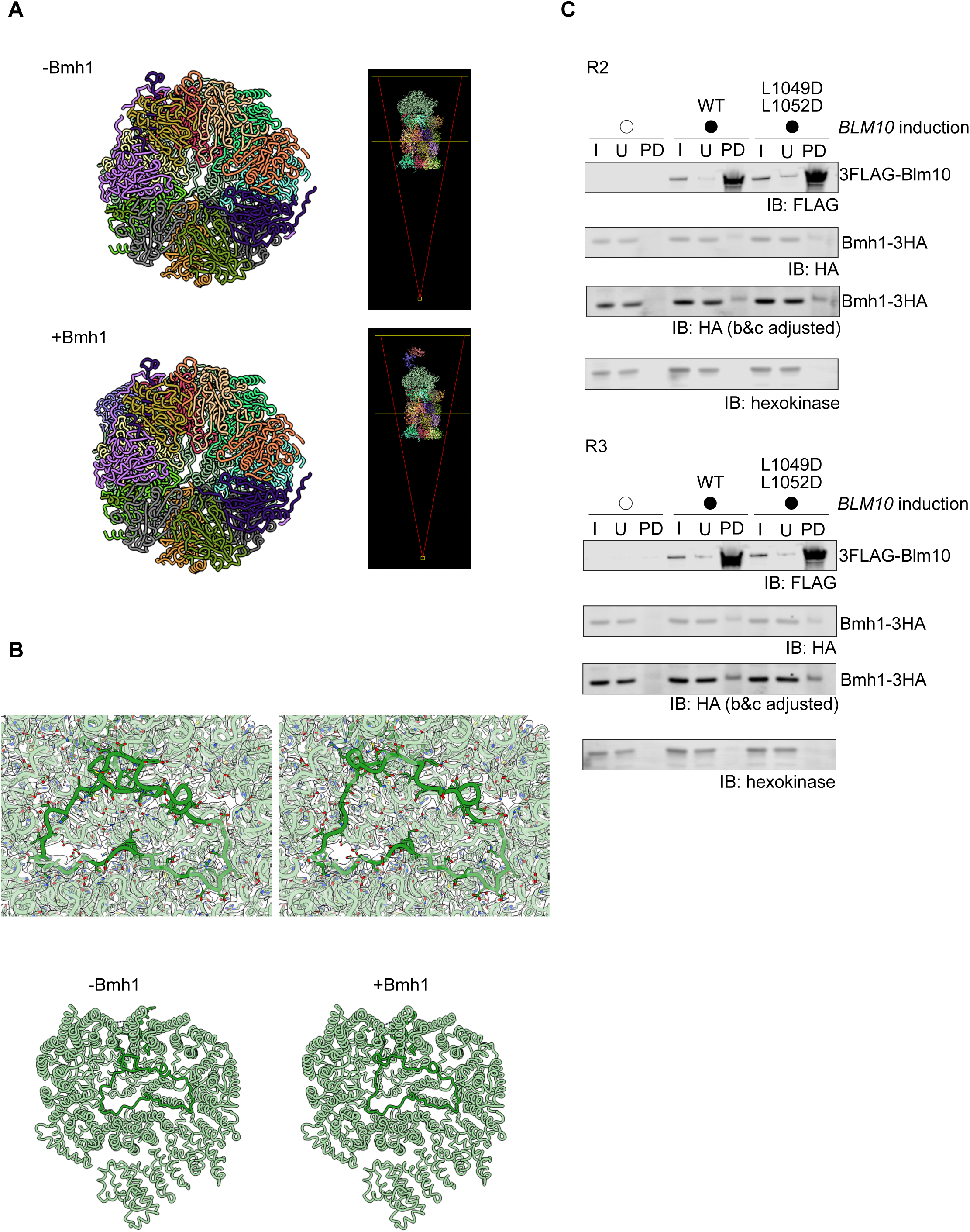
(A) Gate status of the CP without Bmh1 (top) and with Bmh1 bound (bottom), viewed through the internal degradation chamber of the CP (representations to the right). The α-ring is open regardless of Bmh1 binding. (B) Conformational change of a loop within the Blm10 dome that occurs with Bmh1 binding. (C) Second and third replicates (R2 and R3) for co-IP of 3FLAG-Blm10 in mitotic cells. I = input, U = unbound, PD = pulldown.

## Methods

### Yeast strains, plasmids, and primers

All strain genotypes and plasmids used in this study are listed Table 1 and Table 2 in respectively. Oligos used to construct strains are listed in Table 3.

Plasmids were constructed by Gibson Assembly (New England Biolabs E2621S) or site-directed mutagenesis (New England Biolabs E0554S). Plasmids were verified via Sanger sequencing or whole-plasmid sequencing.

All yeast strains were constructed in the strain background SK1. Cassette-integration based manipulations were performed using standard PCR-based methods (Knop et al., 1999; Longtine et al., 1998), with the primers listed in Table S3. Single-copy transgenes were digested with PmeI (New England Biolabs R0560S) and integrated into the genome. Yeast transformation was performed using the lithium acetate method, and integration was confirmed by PCR, Western blot, or fluorescence microscopy when applicable.

### Growth and sporulation conditions

All yeast strains were thawed to YPG (1% yeast extract, 2% peptone, 2% glycerol) plates overnight and patched to YPD (1% yeast extract, 2% peptone, 2% glucose, for haploids) or YPD4% (1% yeast extract, 2% peptone, 4% glucose, for diploids) plates for 8-24 hours prior to starting liquid cultures. Cells were grown in YPD supplemented with tryptophan and uracil at 30°C with shaking for mitotic liquid experiments. All cultures were grown at volumes that were 1/10^th^ of the flask volume to allow for proper aeration.

For meiosis experiments, cells were allowed to saturate overnight in YPD, then diluted into nonfermentable carbon source growth media (BYTA, 1% yeast extract, 2% bacto tryptone, 1% potassium acetate, and 50mM potassium phthalate) at OD_600_ = 0.25 for pre-meiotic starvation for 16 hours at 30°C with shaking. BYTA cultures were then pelleted, washed with either sterile MilliQ H_2_O or sporulation medium (SPO, 2% potassium acetate, 40 mg/L adenine, 40 mg/L uracil, 10 mg/L histidine, 10 mg/L leucine and 10 mg/L tryptophan adjusted to pH 7.0), then resuspended in SPO at OD_600_ = 1.85. SPO cultures were allowed to grow with shaking at 30°C for the duration of the experiments.

For induction of Blm10, either 1 µM beta-estradiol (β-ER) (Sigma Aldrich E8875) or EtOH was added to LexA/LexO strains, or 5 μg/mL anydrotetracycline (aTc) (Cayman Chemical 10009542) or DMSO was added to TetR/TetO strains. For β-ER-based induction, cells were allowed to grow to log phase (around OD_600_ = 0.6) then induced for 1 hour before sample collection. For aTc-based induction, cells were backdiluted, treated with aTc/DMSO, then allowed to grow to log phase (around OD_600_ = 0.6) for collection.

### Meiotic staging

Meiotic staging was performed scoring DAPI morphology (mono-, bi-, and tetra-nucleate cells) by fluorescence microscopy. Samples were fixed in 3.7% formaldehyde overnight at 4°C. Cells were then washed with 0.1M KPi pH 6.4 and resuspended in 20-100 μL of KPi sorbitol pH 7.5. Cells were then mounted in slides coated in poly-L-lysine and washed with 70% EtOH. DAPI + mounting medium (Vectashield H-1200, Vector Labs) was deposited on top of cells and slides were sealed with #1 coverslips and clear nail polish. Imaging conditions for DAPI are found under “Fluorescence microscopy.”

### Measurement of sporulation efficiency, spore viability and colony size

Sporulation of cells was measured after 24 hours in SPO culture under a light microscope. Cells were scored as unsporulated/monads, dyads, or triads/tetrads.

Spore viability was measured after digesting spores incubated on a SPO plate for 24 hours at 30°C with 1 mg/mL zymolyase (MP Biomedicals) for 4 minutes at room temperature, then dissecting spores from individual tetrads onto YPD plates. Plates were incubated for 48 hours 30°C, at which when growth was measured. To assess respiration competency, dissection plates at 48 hours were also replica plated to YPG plates and incubated at 30°C for 24 hours.

For measurement of spore colony size, plates with dissected spores were (originating from SPO plates) were incubated at 30°C for 48 hours post dissection. Plates were photographed using a Geldoc XR+ (Bio-Rad), and colony sizes were quantified on FIJI (2.3.0).

Prism Graphpad (version 10.1.0) was used to plot and analyze data. Statistical significance was assessed using the unpaired t-test.

### Microfluidics timelapse imaging of gamete replicative lifespan

The replicative lifespan of gametes from aged and young progenitors was determined as previously described (Spiri et al., 2025). Briefly, Young and aged cell populations were sorted by biotin-labeling of a founder population followed by bead-mediated magnetic sorting as described Spiri et al. (Smeal et al., 1996). Following sorting immunostaining on young and aged cell suspensions to determine number of bud scars and efficiency of biotin labelling was performed using Wheat Germ Agglutinin, Alexa 350 Fluor conjugate (1 μg/mL, ThermoFisher Scientific) and anti-streptavidin (564 nm) (1 μg/mL). Sporulation cultures were setup as described in (Spiri et al., 2025). Cultures were incubated for 24 hours at 30°C in the dark to complete gametogenesis and post gametogenic age sorted tetrads were separated into single gametes via sonication after 3h incubation of 1 mL cell suspension with zymolyase (1 mg/mL) and 2 μL β-mercaptoethanol as described.

Microfluidics devices (adapted from (Jo et al., 2015); 18++ chips from iBiochips) were setup using the manufacturer’s recommendations on an ECHO Revolution fluorescence microscope in the inverted mode with the environmental chamber set to 30°C. Briefly, 20 mL syringes (Becton, Dickinson and Company) filled with ∼15 mL of YPD 2% + ampicillin (100 μg/mL) + 1% Pluronic F-127 surfactant (P2443, Sigma) were connected to a Clay Adams Intramedic Luer-Stub Adapter (Becton, Dickinson and Company). Non-DEHP Medical Grade Tubing (iBiochips, MTB-100, ID = 0.202”, OD = 0.060”, wall = 0.20”) equipped with a hollow stainless-steel pin (iBiochips, HSP-200) was then used to connect the media syringe to the imaging units using the “media loading” ports. “Cell loading” ports were plugged with solid stainless-steel pins (iBiochips, SSP-200) while imaging units were primed with media for at least 2 hours with a flow rate of 2.4 μL/min using micropumps (iBiochips,78-7200BIO). After all units were filled with media, pumps were stopped an hour prior to cell loading. For cell loading, 5 mL syringes (Becton, Dickinson and Company) with Clay Adams Intramedic Luer-Stub Adapter (Becton, Dickinson and Company) were filled with cells diluted to an OD_600_ of 0.8 in YPD 2% + ampicillin (100 μg/mL). Solid stainless-steel pins were then removed from the “cell loading” ports and tubing and open stainless-steel pins were used to connect the syringe containing cells with the microfluidic device. Cells were loaded manually and, once sufficient cells were loaded, the “cell loading” ports were plugged with solid stainless-steel pins and media flow was resumed at 2.4 μL/min. Timelapse imaging was performed using an Olympus 40x/1.4 NA oil-immersion objective, at 15-20 minute for 72 hours. Image acquisition for chamber was achieved with 2x27 tiled fields of view, with three 3-5 µm z-slices.

After a movie was completed and processed, single Pma2-GFP-positive cells (gametes) were identified and cropped. Next, the replicative lifespan of individual gametes was assessed in the brightfield channel by counting the number of daughter cells produced.

### Quantification of replicative age and purity of sorted cells

Z-stacks (36, 0.2 µm spacing) of fixed young and aged cells/tetrads were acquired on a DeltaVision Elite wide-field fluorescence microscope (GE Healthcare), using a 60x/1.42 NA oil-immersion objective and deconvolved with softWoRx imaging software (GE Healthcare). To reliably count number of bud scars, images were cropped, 3D projected and counted using a custom semi-automated FIJI script (ImageJ2, V. 2.14.0/1.54m, (Schindelin et al., 2012)). Max intensity Z-projections were generated and displayed as examples in figures.

Prism Graphpad (version 10.1.0) was used to plot and analyze data. Prism Graphpad was used to calculate mean and standard deviation of replicative ages and to assess statistical significance in differences between replicative lifespans using the Log-rank (Mantel-cox) test.

### Spot assay

Strains were allowed to saturate overnight in YPD at 30°C with shaking. Cultures were then backdiluted to OD_600_ = 0.2 and 5-fold serial dilutions were performed in a 96 well plate. 2.5 μL of each serial dilution was plated on YPD plates with a multichannel pipette. Plates were then incubated at either 30°C, 40°C (for strains with many prototrophy markers), or 34°C (for strains with no prototrophy markers) for 48 hours. Growth was monitored at 24 hours and 48 hours.

### Growth curve analysis

Strains were allowed to saturate overnight in YPD at 30°C with shaking. Cultures were then backdiluted to OD_600_ = 0.2 in 200 μL of YPD per well, and run in technical triplicates, in a round-bottom 96-well plate (Greiner Cellstar 650 180) OD_600_ was monitored, with shaking at 30°C, for about 20 hours using a plate reader (either a Versa Max Plate Reader (Molecular Devices) or Spark Plate Reader (Tecan)). Haploid cells were sonicated 3 times at 50% to minimize clumping. Doubling time was determined with the slope from the linear portion of the growth curve.

### Fluorescence microscopy and timelapse imaging of germination

All fluorescence microscopy was performed using a DeltaVision Elite wide-field microscope (GE Healthcare), using a 100x/1.40 NA oil-immersion objective and a PCO Edge sCMOS camera. Cells were fixed in 3.7% formaldehyde overnight at 4°C. Cells were then washed with 0.1M KPi pH 6.4 and resuspended in 20-100 μL of KPi sorbitol pH 7.5. Cells were imaged with the following settings:

Hsp104-eGFP: 32%T, 0.05s, EX: 475/28 EM: 523/36.

Htb1-mCherry: 32%T, 0.05s, EX: 575/25 EM: 632/60.

DAPI: 100% T, 0.1s, EX: 390/18 EM: 435/48

For germination movies, cells from 24 hours in SPO culture were washed with MilliQ H_2_O twice, then resuspended in MilliQ H_2_O and sonicated 3 times at 50% to separate out cell clumps. Cells were then deposited on concanavalin-A-treated glass-bottom wells (Corning 4580) 100 μL of YPD was added to each well. Timelapse imaging was performed on a DeltaVision Elite microscope (GE Healthcare) for 9 hours, with POL images taken every 5 minutes, with the environmental chamber set to 30°C.

Images were deconvolved using the softWoRx imaging software (GE Life Sciences) and z-projected with FIJI.

Prism Graphpad (version 10.1.0) was used to plot and analyze data. Statistical significance was assessed using the unpaired t-test.

### Mass spectrometry

For meiotic sample collection, 250 mL of SPO culture at OD_600_ = 1.85 were harvested per timepoint by filtration and flash freezing in liquid nitrogen with lysis buffer with protease inhibitors (20 mM Tris-HCl pH 7.5, 140 mM KCl, 1.5 mM MgCl_2_, 1% Triton-X 100, 2 µg/mL aprotinin, 10 µg/mL leupeptin, 1mM PMSF, 1:100 PIC2 (Sigma), 1:100 PIC3 (Sigma), 1x Ultra cOmplete protease inhibitor cocktail (Roche), 1x Pepstatin A (Sigma), For mitotic samples, 750 mL of YPD culture at log phase (OD_600_ = ∼0.6) were harvested. Samples were lysed by Retsch mixermilling (6x 3-minute rounds at 15 Hz). Resulting powder was thawed, spun once at 4C for 5 min at 3000g, sup was removed and spun at 20,000g at 4C for 10 minutes. Extract was aliquoted in 200ul portions and flash frozen. Identical extract was used for mass spectrometry and mRNA-sequencing.

For sample processing for LC-MS/MS measurements of the global proteome of WT and *blm10*Δ cells, Proteins were precipitated and desalted using the SP3 method, as described in (Hughes et al., 2019). Samples were lysed in 8M Urea buffer (8M Urea, 75mM NaCl, 50mM Tris/HCl pH 8.0, 1mM EDTA) and protein concentration using the Pierce™ BCA protein assay kit (Thermo Scientific) based on the manufacturer’s instructions. 40 µg from each lysate were processed. Disulfide bonds were reduced with 5 mM dithiothreitol and cysteines were subsequently alkylated with 10 mM iodoacetamide. Proteins were precipitated on 0.5 µg/µL speedBead magnetic carboxylated modified beads (1:1 mix of hydrophobic and hydrophilic beads, cat# 6515215050250, 45152105050250, GE) by addition of 100% ethanol in a 1:1 (vol:vol) sample:ethanol ratio followed by 15 min incubation at 25°C, 1000 rpm. Protein-bound beads were washed in 80% ethanol and proteins were digested off the beads by addition of 0.8 µg sequencing grade modified trypsin (Promega) in 100 mM Ammonium bicarbonate, incubated 16 hrs at 25°C, 600 rpm. Beads were removed and the resulting tryptic peptides evaporated to dryness in a vacuum concentrator. Dried peptides were then reconstituted in 15 μl of 3% ACN / 0.2% Formic acid.

For sample processing for LC-MS/MS measurements of aggregates, proteins were precipitated as follows: 20 µL of 100% trichloroacetic acid (TCA) was added to 200 µL of lysate, and samples were incubated at 4°C for 30 minutes. Samples were then centrifuged at 14,000 rpm for 5 minutes at 4°C, and the supernatant was removed. Pellets were washed with cold acetone to remove residual TCA, followed by centrifugation at 18,000 × g for 5 minutes at 4°C. The resulting pellets were air-dried for 1.5 minutes to allow remaining acetone to evaporate and subsequently resuspended in 50 µL of 8 M urea for downstream processing.

A total of 40 μg of protein from protein precipitation, quantified using the Pierce™ BCA Protein Assay Kit (Thermo Scientific) according to the manufacturer’s instructions, was reduced with 5 mM dithiothreitol (DTT) for 45 minutes at 25 °C with shaking at 600 rpm. Reduction was followed by alkylation with 10 mM iodoacetamide (IAA) for 45 minutes in the dark under the same conditions. Samples were then diluted with 50 mM Tris-HCl (pH 8.0; Thermo Scientific) to reduce the urea concentration to below 2 M. Proteins were digested overnight at 25 °C and 600 rpm using 0.8 μg of sequencing-grade modified trypsin (Promega). After digestion, peptides were acidified with formic acid (Thermo Scientific) and desalted using in-house packed C18 StageTips (two plugs), following the protocol described by (Rappsilber et al., 2007). Cleaned peptides were dried using a Thermo Savant SpeedVac and reconstituted in 3% acetonitrile/0.2% formic acid prior to LC-MS/MS analysis.

For LC-MS/MS analysis on a Q-Exactive HF, about 1 μg of total peptides were analyzed on a Waters M-Class UPLC using a IonOpticks Aurora ultimate column (1.6 µm, 75 µm x 25 cm for global proteomes, 1.7 µm, 75 µm x 15 cm for aggregate measurements) coupled to a benchtop Thermo Fisher Scientific Orbitrap Q Exactive HF mass spectrometer. Peptides were separated at a 400 nL/min flow rate with a 100-minutes and 90-minute gradient, (for global measurement and aggregate measurements, respectively), including sample loading and column equilibration times, using solvents A (0.1% formic acid in water) and B (0.1% formic acid in acetonitrile). The detailed gradients of solvent B are: 2% B for 1 min; linear increase to 10% B over 29 min; linear increase to 22% B over 27 min; linear increase to 30% B over 5 min; linear increase to 60% B over 4 min; linear increase to 90% B over 1 min, held for 2 min; linear decrease to 50% B over 1 min, held for 5 min; linear decrease to 2% B over 1 min; and re-equilibrated at 2% B for the rest of the acquisition period. Data were acquired in data-independent mode using Xcalibur software (4.5.474.0). MS1 spectra were measured with a resolution of 120,000, an AGC target of 3e6, and a scan range from 350 to 1600 m/z. For global proteome measurements, MS2 spectra were measured in 34 segment windows per MS1, each with an isolation window width of 38 m/z (0.5 m/z overlap with the neighboring window), a resolution of 30,000, an AGC target of 3e6, and a stepped collision energy of 22.5, 25, 27.5. For aggregate measurements, 21 isolation windows of 60 m/z were measured at a resolution of 30,000, an AGC target of 3e6, normalized collision energies of 22.5, 25, 27.5, and a fixed first mass of 200 m/z.

For data analysis of global proteome measurement, all raw data were analyzed with SpectroNaut software version 16.0.220606.53000 (Biognosys) using a directDIA method based on a UniProt yeast database (release 2014_09, Saccharomyces cerevisiae strain ATCC 204508 / S288c), performed with the “BGS factory settings” including the following parameters: Oxidation of methionine and protein N-terminal acetylation as variable modifications; carbamidomethylation as fixed modification; Trypsin/P as the digestion enzyme; For identification, we applied a maximum FDR of 1% separately on protein and peptide level. “Cross run normalization” and “Global imputing” were activated. This gave intensity values for a total of 4358 protein groups across all samples and replicates. “PG.Quantity” (normalized across samples) values were used for all subsequent analyses.

For data analysis of aggregate measurements, raw DIA data were processed using DIA-NN (version 2.1.0) (Demichev et al., 2020). DIA-NN was used in library-free mode, in which an in silico spectral library was generated directly from the *Saccharomyces cerevisiae* protein sequence database (UniProt proteome: UP000002311). The predicted spectral library was then used for peptide identification and protein quantification across all samples. The analysis were performed using default DIA-NN settings, with protein N-terminal acetylation as variable modification, match between runs enabled and protein inference enabled.

For downstream analysis of global proteome and aggregate measurements, protein abundance values from each sample were log-adjusted and normalized in Cluster 3.0 prior to clustering. Hierarchical clustering was performed on genes based on a centered correlation-based similarity metric and centroid linkage. Hierarchical clustering was visualized with Java Treeview (1.2.0, (Saldanha, 2004)). GO term analysis visualizations were done with R packages ggplot2 (4.0.1, (Wickham, 2016)) and ClusterProfiler (4.18.2, (Yu et al., 2012)). Differential analysis of MS abundance for global proteome and aggregate measurements was done with R packages DEqMS (1.28.0, (Zhu et al., 2020)) and Limma (3.66.0, (Ritchie et al., 2015)).

### Two-phase fractionation for analysis of protein aggregation

Two-phase fractionation was adapted from two previous protocols (Hamdan et al., 2017; Jang et al., 2004). Approximately 15 ODs of cells were harvested after growing to log phase (OD_600_ = ∼0.6). Cells were washed in spheroplast buffer (1x PBS, 1M sorbitol, 1mM EDTA) resuspended in cold lysis buffer with protease inhibitors (50 mM potassium phosphate, pH 7.0, 1mM EDTA, 5% glycerol; 1 mM PMSF, 1x Ultra cOmplete EDTA-free protease inhibitor cocktail (Roche 11873580001)) then flash frozen in liquid nitrogen. Cell solutions were thawed on ice and bead beaten with 0.5mm zirconia beads (BioSpec) for 5 minutes for a total of 3 cycles, with resting on ice for 2 minutes in between. Beads were washed with lysis buffer to collect residual lysate, then samples were spun at 800g for 10 minutes to remove intact cells. Supernatant was transferred to a new tube, then spun at 13,000g for 20 minutes at 4°C. Supernatant was removed (“soluble”/ “S” fraction). 3x SDS sample buffer (250 mM Tris-HCl, pH 6.8, 8% BME, 40% glycerol, 12% SDS, 0.00067% bromophenol blue) was added to soluble fractions to a final concentration of 1x. Pellets were resuspended in lysis buffer + NP-40, centrifuged at 13,000g for 20 min at 4°C, and supernatant was removed. Pellet resuspension and high-speed spins were repeated once more. Pellets were resuspended in 1x SDS/urea sample buffer (3x SDS sample buffer diluted to 1x, with 8M urea). Aggregate samples were spun at 20,000g for 5 minutes at room temperature and the supernatant (“aggregate”/ “A” fraction).

### RNA extraction, library preparation, and mRNA sequencing

Identical extract was used for mass spectrometry and mRNA-sequencing (see sample collection in “Mass spectrometry”).

An equal volume of citrate-buffered acid phenol (pH 4.3, P4682, Sigma-Aldrich) was added to whole cell extracts and incubated at 65°C for 30 min in a Thermomixer (Eppendorf) shaking at 1400 rpm. After microcentrifugation (20,000g for 10 min at 4°C) the aqueous phase was transferred to a second tube with 350 μl chloroform. The aqueous phase was again separated by microcentrifugation (20,000g for 5 min at room temperature) and RNA was precipitated in 100% isopropanol with 350 mM sodium acetate (pH 5.2) overnight at –20°C. Pellets were washed with 80% ethanol and resuspended in DEPC water. Total RNA was quantified by Nanodrop. 5 μg of total RNA was used for mRNA isolation with NEXTflex Poly(A) Beads 2.0 (NOVA-512991, Revvity).

RNA-seq libraries were generated with the NEXTflex Rapid Directional RNA-Seq Kit 2.0 (NOVA-5198-02, Revvity) according to manufacturer’s instructions. AMPure XP beads (A63881, Beckman Coulter) were used to select fragments between 200-500 bp. Libraries were quantified using the Agilent4200 TapeStation (Agilent Technologies, Inc). Single-read sequencing was performed at the UCSF CAT, supported by UCSF PBBR, RRP IMIA, and NIH 1S10OD028511-01 grants, with a NovaSeqX.

Hisat2 (v2.1.0, (Kim et al., 2019)) was used to align reads to the sacCer3 reference genome (v64). Quantification of RNA as transcripts per million (TPM) was done with StringTie (v2.1.6, (Pertea et al., 2015)). Hierarchical clustering was performed using Cluster 3.0 (de Hoon et al., 2004) and visualized in Treeview (v1.2.0, (Saldanha, 2004)). TPM values of each sample were log-adjusted and normalized in Cluster 3.0 prior to clustering. Hierarchical clustering was performed on genes based on a centered correlation-based similarity metric and centroid linkage.

### Native PAGE and in-gel peptidase assay

For meiotic and mitotic samples, 25 mL of SPO or 50 mL log phase culture (∼30-46 ODs) was harvested. For SPO cultures, 2 mM PMSF was added to media immediately prior to harvest. Cells were pelleted and flash frozen in liquid nitrogen. 250 mL of lysis buffer (60 mM HEPES pH 7.6, 25 mM NaCl, 10 mM MgCl_2_, 2.5% glycerol, 0.25% Triton-X 100, PhosSTOP tablets (2 tablets used per 20 mL buffer, Sigma), Ultra cOmplete protease inhibitor tablet (1/2 tablet used per 20 mL buffer, Sigma), 0.64 µg/mL AEBSF (Sigma), 2.7 µM Pepstatin A (Sigma)) supplemented with an ATP regeneration system (5 mM ATP (Sigma Aldrich A6419), 16 mM creatine phosphate (VWR 0271), 0.05 mg/mL creatine kinase (Sigma Aldrich C3755)) and 250 mL 0.5mm zirconia beads (BioSpec) was added to thawed cell pellets. Samples were lysed on a FastPrep-24 (MP Biomedicals) for 1 round at 40 seconds, 0.6 m/s. Lysates were cleared by low-speed centrifugation (3000 rpm for 20 seconds), followed by high-speed centrifugation (20,000g for 1 minute). Protein content was quantified by Bradford or BCA assay (Bio-Rad 5000006, Thermo Scientific 23225). Absorbances were measured by a Versa Max Plate Reader (Molecular Devices). A portion of the lysate for each sample was diluted to 1 mg/mL with GF buffer (60 mM HEPES, pH 7.6, 25 mM NaCl, 25 mM KCl, 5 mM MgCl_2_, 0.5 mM EDTA, 5% Glycerol) supplemented with 5 mM ATP and 0.5 mM DTT.

Native PAGE and the in-gel peptidase assay was performed as described in (Roelofs et al., 2018). Specifically, 10-well 3.5% polyacrylamide (National Diagnostics Protogel 37.5:1 acrylamide:bis-acrylamide, EC-890) gels were poured with 1x native gel running buffer (90 mM Tris, 90 mM boric acid, 5 mM MgCl_2_, 0.5 mM EDTA) and supplemented with 0.5 mM ATP. 10 μL of 1 mg/mL lysate with 5x native loading buffer (250 mM Tris-HCl pH 7.5, 50% glycerol (v/v), ∼50 ng/mL xylene cyanol) used at a final concentration of 1x was loaded per well. Gels were run at 100V for 3 hours at 4°C in native gel running buffer (90 mM Tris, 90 mM boric acid, 5 mM MgCl_2_, 0.5 mM EDTA, 0.5 mM ATP). To visualize proteasome activity, gels were incubated for 15 minutes at 30°C in an activity assay buffer (0.05M Tris-HCl, 5 mM MgCl_2_, 1 mM ATP) containing 500 µM suc-LLVY-amc (Bachem), imaged once on a Geldoc XR+ (Bio-Rad) with UV transillumination at 302 nm (ethidium bromide settings), incubated with 0.02% SDS for another 15 minutes at 30°C to visualize inhibited proteasomes, then imaged again.

### Co-immunoprecipitation

Sample collection and cell lysis was performed as in “Native PAGE and in-gel peptidase assay.” 50-100 μL of epitope affinity matrix (anti-HA affinity matrix (Roche 11 815 016 001), anti-V5 affinity matrix (Sigma Aldrich A7345)) were washed with GF buffer (60 mM HEPES, pH 7.6, 25 mM NaCl, 25 mM KCl, 5 mM MgCl_2_, 0.5 mM EDTA, 5% Glycerol and 0.5mM DTT), then 100-200 μL of 1 mg/mL lysate was applied. The same amount of lysate was used for each sample for each experiment. Input (“I”) samples were collected prior to application to beads. Lysates and beads were incubated with rotation at 4°C for 1 hour. Unbound (“U”) samples were collected from the supernatant. Beads were then washed 3 times with GF buffer with 0.5 mM ATP.

Beads were resuspended in 20 μL of GF buffer. 3x SDS sample buffer (250 mM Tris pH 6.8, 8% β-mercaptoethanol, 40% glycerol, 12% SDS, 0.00067% bromophenol blue) was added to samples to a final concentration of 1x, then boiled at 95°C for 5 minutes to elute bound proteins and spun for 5 minutes at 20,000g (pulldown/ “PD” samples).

### Western Blotting

About 5 ODs of mitotic or meiotic culture were pelleted and resuspended in cold 5% tricholoracetic acid (TCA) and incubated at 4°C for at least 15 minutes. Pellets were washed with acetone and allowed to air dry overnight. Pellets were then combined with 100 μL protein breakage buffer (50 mM Tris pH 7.5, 1 mM EDTA, 3 mM DTT, 1X cOmplete EDTA-free inhibitor cocktail (Roche 11873580001), 2 mM PMSF) and 100 μL acid-washed glass beads (Sigma Aldrich G8772), then bead-beat for 5 minutes on a Mini-Beadbeater-96 (BioSpec). 50 µL of 3x SDS sample buffer (250 mM Tris pH 6.8, 8% β-mercaptoethanol, 40% glycerol, 12% SDS, 0.00067% bromophenol blue) was added to the samples.

Samples were boiled at 95°C for 5 minutes and spun at 20,000g for 5 minutes prior to loading on 4-12% Bis-Tris Bolt gels (Invitrogen NW04125BOX). Gels were run at 110V for 1.5 hours in 1x MES buffer (Invitrogen B000202). 2 μL of TCA extract was loaded per sample, along with 3 μL of PageRuler Plus Protein ladder (Thermo Scientific 26620). Wet transfer to 0.2µm PVDF membranes (Bio-Rad 1620177) were performed with a Mini-PROTEAN Tetra tank (Bio-Rad) in 9% Towbin Buffer (25 mM Tris, 192 mM glycine, 9% MeOH), run at a 180 mA constant, 80V max for 3 hours at 4°C.

Membranes were blocked in PBS Intercept Blocking Buffer (LI-COR Biosciences). Blots were incubated overnight in primary antibody diluted in PBS Intercept Blocking Buffer +0.1% Tween-20 at 4°C. The following primary antibodies were used:

1:1,000 rabbit anti-Blm10 (Enzo BML-PW0570-0025)

1:1,000 mouse anti-HA.11 (Biolegend 901515)

1:1,000 rabbit anti FLAG (Cell Signaling Technology 23685)

1:1,000 mouse anti-GFP (JL-8) (Clontech)

1:1,000 mouse anti-3V5 (Invitrogen)

1:10,000 rabbit anti-hexokinase (Rockland)

1:10,000 rat anti-tubulin (Serotec)

Blots were then washed in PBST (PBS + 0.1% Tween-20) and incubated in secondary antibody diluted in PBS Intercept Blocking Buffer + 0.01% Tween-20 for 1 hour at RT. The following secondary antibodies were used:

1:15,000 goat anti-mouse conjugated to IRDye 800CW (LI-COR Biosciences)

1:15,000 donkey anti-mouse conjugated to IRDye 800CW (LI-COR Biosciences)

1:15,000 goat anti-rabbit conjugated to IRDye 680 RD (LI-COR Biosciences)

1:15,000 donkey anti-rabbit conjugated to IRDye 680 RD (LI-COR Biosciences)

Blots were then washed in PBST and imaged using the Odyssey CLx system (LI-COR Biosciences). Image analysis and quantification was performed in ImageStudioLite 5.2.5.

### Protein purification

*S. cerevisiae* Blm10 and Blm10-20S core complexes were purified from a yeast strain overexpressing 3x-Flag-*BLM10* under a Tet-inducible system ((Azizoglu et al., 2021), ÜB 42752, Table 1). Strains were grown to log phase (OD_600_ = ∼0.6), then induced with 5 μg/mL aTc for 3-4 hours, then harvested and flash frozen with GF buffer (30mM HEPES, pH 7.6, 50mM NaCl, 50mM KCl, 10mM MgCl2, 5% Glycerol and 0.5mM TCEP). Yeast cells were lysed with a 6875 Freezer Mill Dual Chamber Cryogenic grinder (SPEX Sample Prep) and resuspended in lysis buffer (30mM HEPES, pH 7.6, 50mM NaCl, 50mM KCl, 10mM MgCl2, 5% Glycerol and 0.2% NP-40) with an ATP regeneration system (5 mM ATP, 16 mM creatine phosphate, 0.05 mg/mL creatine kinase). The lysate was clarified by centrifugation at 26,000g for 40 minutes and then transferred to 50mL Centrifuge tubes with equilibrated M2 anti-Flag affinity resin (Sigma) and incubated for 1 hour at 4° C. The resin was recovered by centrifugation at 1000g for 10 minutes, washed and transferred to a gravity column. Additionally, the resin was washed with buffer containing 500mM NaCl and 500mM KCl to dissociate 20S core from 3xFlag-Blm10. This step was omitted when purifying Blm10-20S core complexes. The resin was incubated with 3mg/mL 3xFlag peptide-containing buffer to elute the purified protein which was then loaded onto a Superose 6 Increase 10/300 column (Cytiva) equilibrated with GF buffer (30mM HEPES, pH 7.6, 50mM NaCl, 50mM KCl, 10mM MgCl_2_, 5% Glycerol and 0.5mM TCEP).

### Peptidase activity assays

Cleavage of the Suc-LLVY-AMC substrate was monitored by measuring the increase in AMC Fluorescence over time using a BMG Labtech CLARIOstar plate reader at 30° C. Peptidase activity for purified 26S proteasomes, Blm10-20S core and 20S core complexes was determined by combining 25nM of each complex with 50µM Suc-LLVY-AMC substrate and determining the initial rate of substrate cleavage. Blm10 titration experiments were conducted in a similar fashion by incubating 20S core with increasing concentrations of Blm10 and then combining it with Suc-LLVY-AMC to final concentrations of 25nM complex and 50uM substrate. The initial rates of Suc-LLVY-AMC cleavage at varying concentrations of Blm10 were fit to a One site – Specific binding equation using GraphPad (Prism) to calculate a K_D_.

### ATPase stimulation assay

ATP hydrolysis rates were determined using an NADH-coupled assay. 5mM ATP, 3 U ml−1 pyruvate kinase (Sigma), 3 U ml−1 lactate dehydrogenase (Sigma), 1 mM NADH and 7.5 mM phosphoenolpyruvate were mixed with 100nM 26S proteasomes and increasing concentrations of Blm10. NADH absorbance was monitored over time using a BMG Labtech CLARIOstar plate reader at 30° C and the initial rate of NADH depletion was used to calculate ATPase rates. The initial rates of ATP hydrolysis at varying concentrations of Blm10 were fit to a Dose response -Inhibition equation using GraphPad (prism) to determine an IC50.

### Cryo-EM sample preparation, data collection, data processing and model building

Blm10-20S core complexes were diluted in GF buffer (30mM HEPES, pH 7.6, 50mM NaCl, 50mM KCl, 10mM MgCl2, 5% Glycerol and 0.5mM TCEP) with 0.02% NP-40 to a final concentration of 4mg/mL. 3uL microliters of the sample were pipetted on a glow-discharged (25mA for 25s) UltrAufoil R 2/2 200-mesh gold grid (Quantifoil), blotted for 2.5s and immediately plunge-frozen in ethane using a Vitrobot Mark IV (Thermo Fisher) system set to 12° C. The clipped grid was loaded onto a Titan Krios G2 transmission EM instrument operated at 300 keV (Thermo Fisher) equipped with a Gatan K3 Direct Electron Detector and BIO Quantum Energy Filter. SerialEM was used for automated data acquisition using a physical pixel size of 0.9432 Å (81k magnification) in super-resolution mode, 50-frame movies, a dose of 50 e/Å^2^ and a defocus range of -0.5 to -2μm.

15,116 exposures were pre-processed using CryoSPARC (version 4.6). Blm10-CP particles were picked, extracted, symmetry expanded (C_2_), classified and aligned to reconstruct high-resolution EM volumes. Initial atomic models were generated using ModelAngelo and manual building was carried out using Coot and Isolde followed by real-space refinement in PHENIX. This process was repeated iteratively as necessary.

### Proteasome complex isolation for MS analysis

Cultures were backdiluted after overnight saturation in YPD to OD600 = 0.15 in 500 mL YPD and 5 μg/mL aTc was added. Cells were grown until log phase (OD = 0.6/mL) and were then treated with either 50 μM MG132 and 20 μM carfilzomib (CFZ), or DMSO, for 1 hour. Cells were then resuspended in GF buffer, supplemented with 50 μM MG132 for treated samples, and flash frozen in liquid nitrogen.

Anti-FLAG pulldowns were performed as in “Protein purification” to isolate Blm10-containing complexes. A secondary Ni-NTA pulldown was performed on the anti-FLAG eluate to isolate RP-CP-Blm10 complexes in the eluate fraction and Blm10-containing complexes in the unbound fraction. All buffers for treated samples were supplemented with 50 μM MG132.

## Data availability

All the strains, plasmids, and reagents used in this study are available upon request. Mass spectrometry data and mRNA-sequencing data generated in this study will be made publicly available.

